# Attractor dynamics gate cortical information flow during decision-making

**DOI:** 10.1101/2019.12.14.876425

**Authors:** Arseny Finkelstein, Lorenzo Fontolan, Michael N. Economo, Nuo Li, Sandro Romani, Karel Svoboda

## Abstract

Decisions about future actions are held in memory until enacted, making them vulnerable to distractors. The neural mechanisms controlling decision robustness to distractors remain unknown. We trained mice to report optogenetic stimulation of somatosensory cortex, with a delay separating sensation and action. Distracting stimuli influenced behavior less when delivered later during delay – demonstrating temporal gating of sensory information flow. Gating occurred even though distractor-evoked activity percolated through the cortex without attenuation. Instead, choice-related dynamics in frontal cortex became progressively robust to distractors as time passed. Reverse-engineering of neural networks trained to reproduce frontal-cortex activity revealed that chosen actions were stabilized via attractor dynamics, which gated out distracting stimuli. Our results reveal a dynamic gating mechanism that operates by controlling the degree of commitment to a chosen course of action.

**One Sentence Summary:** Mechanisms controlling state-dependent communication between brain regions allow for robust action-selection.

## Introduction

Transformation of sensory inputs into motor outputs is a fundamental component of decision-making. When sensation and action are separated in time, perceptual decisions about future actions should persist in memory until movement execution. On the neural level this is manifested by preparatory activity observed in frontal cortex and related areas (*1*–*6*) that can predict the upcoming movement (*7*–*14*). During the temporal gap between sensation and action, decisions about upcoming movements (‘motor plans’) can be affected by additional inputs. The mechanisms underlying the robustness of a motor plan to distracting sensory inputs are unknown.

It has been proposed that information flow between sensory and motor areas can be regulated by ‘gating’ mechanisms able to filter out undesirable sensory inputs during stimulus selection (*15*–*21*). Here we set to understand how information flow is regulated during motor-preparation (the delay epoch), by studying the ability of calibrated stimuli to perturb the motor plan for the upcoming movement. We found that inputs of identical strength became progressively less capable of modifying the selected motor plan, which was stabilized by attractor dynamics. Our results identify a novel mechanism that gates information flow between brain areas during motor planning.

## Results

Anterior lateral motor cortex (ALM) is a critical node in a decision-making circuit that controls directional-licking in response to sensory inputs (*8, 11, 13, 14, 22, 23*). To study sensorimotor transformations in this circuit, we developed a delayed-response task, in which mice learned to respond by directional licking to optogenetic stimulation of genetically defined neurons in layer 4 of vibrissal somatosensory cortex (vS1, **Fig. 1A**). A photostimulus applied to left vS1 during the sample epoch (delineated by auditory cues) instructed mice to lick right, whereas the absence of a photostimulus instructed mice to lick left (**Fig. 1B-C**). Time of action was determined by an auditory go cue, played after a 2 s delay epoch. Training mice to respond to photostimulation of sensory cortex (*24*–*26*) allowed us to use calibrated photostimuli and investigate cortical gating mechanisms during decision-making.

**Figure 1.**
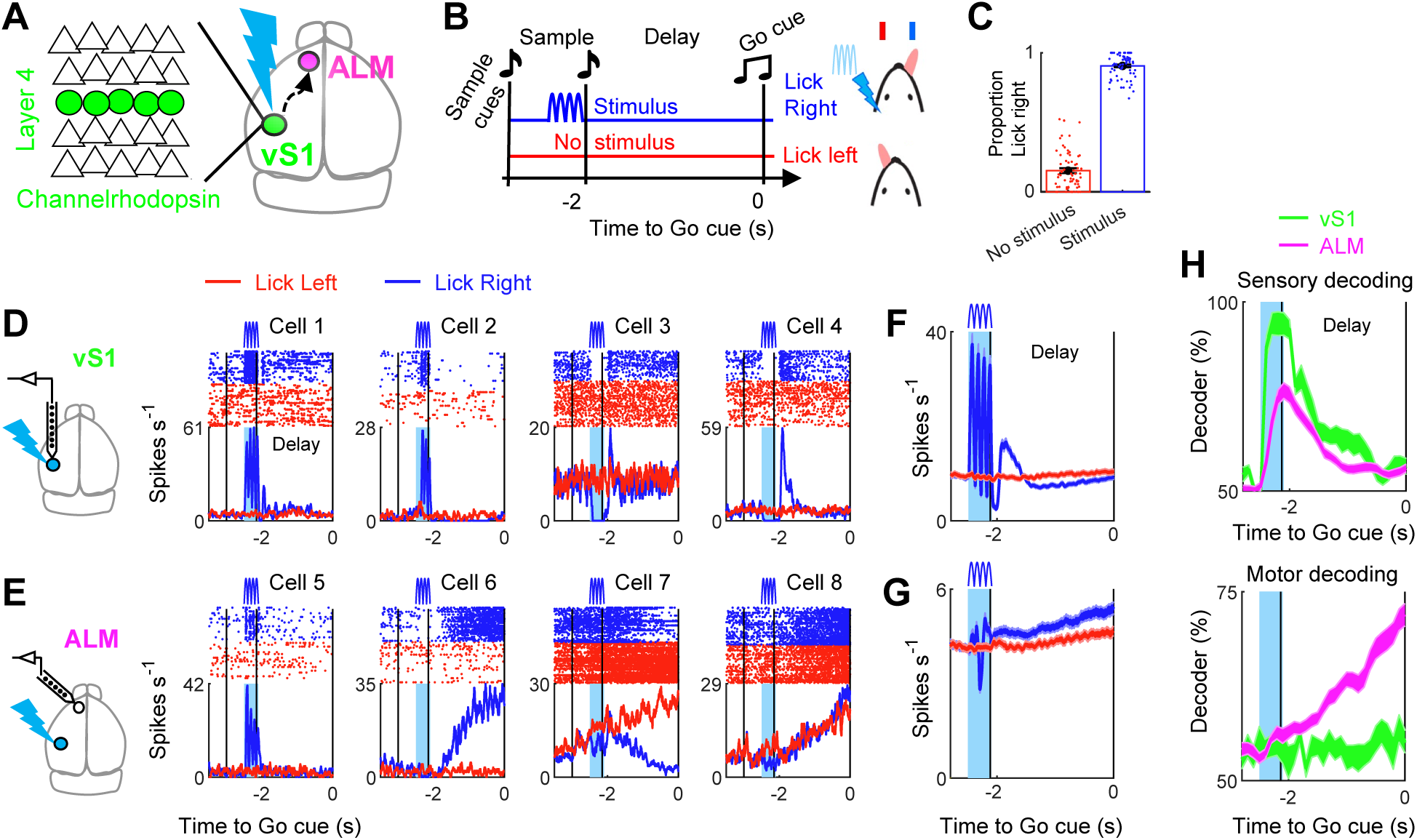
Sensorimotor transformations evoked by direct cortical photostimulation. **A,** Cortical regions involved in tactile decision-making. Left, Schematics of channelrhodopsin-2/EYFP expressing neurons in layer 4 of vS1 (‘barrel cortex’) of transgenic mice. **B,** Basic task. Mice had to lick right in response to vS1 photostimulation during sample epoch (delineated by auditory cues). Absence of stimulus instructed to lick left. A Go cue after a 2 s delay epoch signaled the start of the response epoch. **C,** Behavioral performance (n= 5 mice, 67 sessions); each point corresponds to a behavioral session; mean ± s.e.m. **D-E,** Silicon probes recordings showing example neurons with transient responses in vS1 (D) and a mixture of transient and sustained responses in ALM (E). Spike raster and trial-averaged spike rates on correct lick-right (blue) and lick-left (red) trials are shown for each example neuron. Cyan vertical bar indicates photostimulation. Black vertical bars delineate trial epochs. **F-G,** Grand-average population responses of putative pyramidal cells on lick-right (blue) and lick-left (red) trials in left vS1 (**F**) and left ALM (**G**). **H,** Decoding sensory-related (top) and choice-related (bottom) activity in vS1 and ALM, at different times along the trial.

We used silicon probes to record activity from individual putative pyramidal cells in vS1 (the site of photostimulation) and ALM (**Fig. 1D-E, Sup. Fig. 1;** for putative fast-spiking interneurons see **Sup. Fig. 2**). Photostimulation transiently modulated the spike rate of most (58%) vS1 neurons (**Fig. 1D**). On the population level, response of vS1 neurons peaked around the time of photostimulation and decayed rapidly thereafter (**Fig. 1F**). In ALM, only a small proportion (13%) of neurons responded transiently to photostimulation (**Fig. 1E**, example cell 5). These neurons were more abundant on the side of the photostimulated vS1 (left hemisphere; **Sup. Fig. 1B**, cluster 4). The majority of ALM neurons showed preparatory activity during the delay epoch (**Sup. Fig. 1B**, clusters 1-3,6), as trial type selective (**Fig. 1E**, Cell 6-7) or non-selective (**Fig. 1E**, Cell 8) ramping.

**Figure 2.**
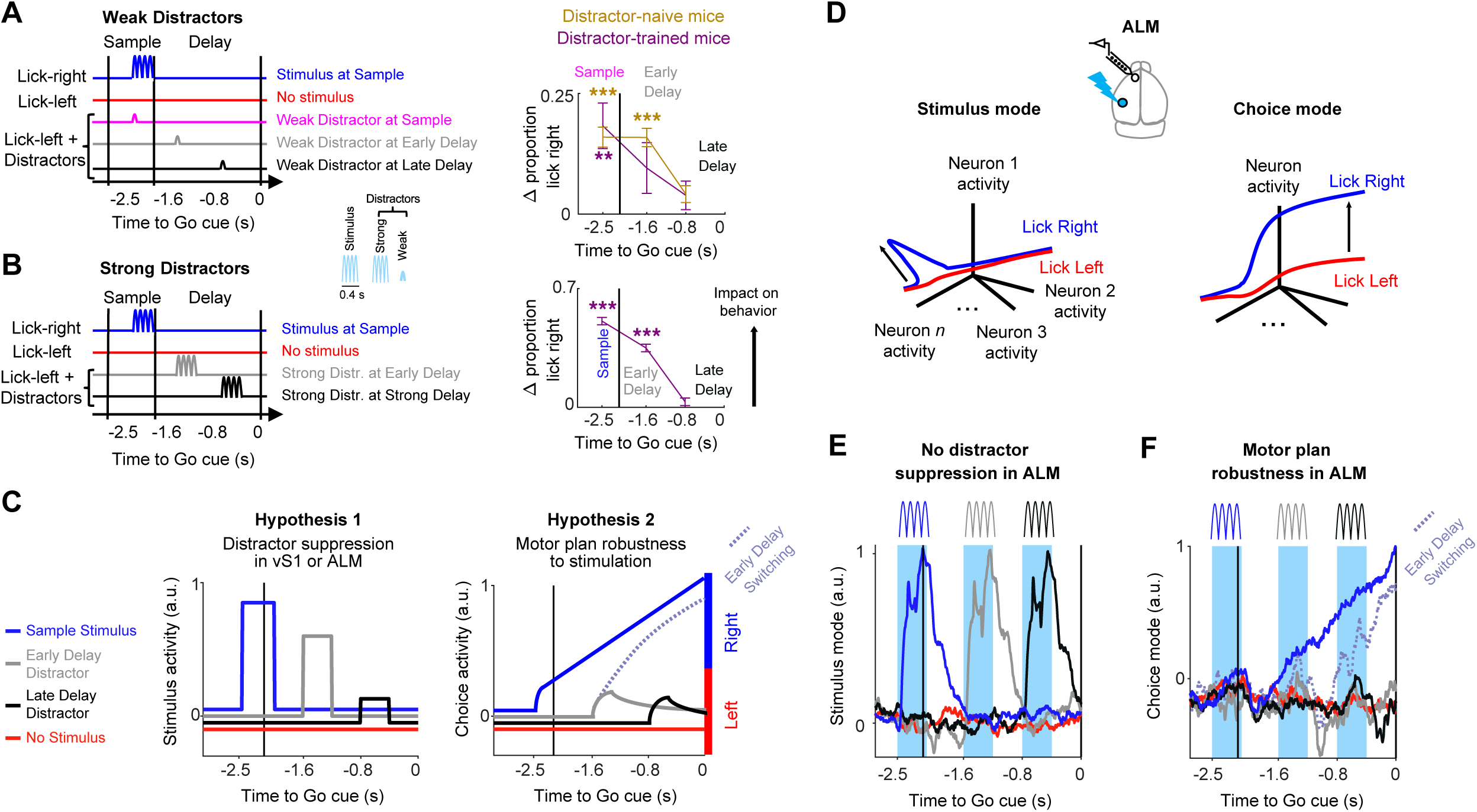
Temporal gating of photostimuli by population dynamics in ALM. **A-B**, Left, task schematics with weak (A) and strong (B) distractors. Right, behavioral impact of distractors delivered at different times in distractor-naive (gold) and distractor-trained mice (purple). Behavioral impact was quantified as change in proportion of lick-right responses on photostimulation trials, relative to no-stimulus trials. Error bars, mean ± s.e.m., across sessions; * *P* < 0.05, *** *P* < 0.001 by paired Student t-test, photostimulation trials versus non-stimulus trials. **C,** Possible mechanisms for temporal gating. **D,** Targeted dimensionality reduction approach to define Stimulus and Choice modes in neural activity space. **E-F**, Session-averaged neural activity of left ALM projected on Stimulus (**E**, all trials) and Choice (**F**, correct trials only) modes in response to vS1 photostimulation. Lick-left trajectory without stimulation (red), with distractors during early-delay (gray), or late-delay (black). Lick-right trajectory during sample-epoch stimulation (blue). Black bars delineate the delay epoch. Dashed-gray, trials with early-delay distractors that resulted in lick-right response (‘switching trials’). Timing of photostimulation on different trial types is indicated by stimulus profile icons and cyan vertical bars.

**Figure 3.**
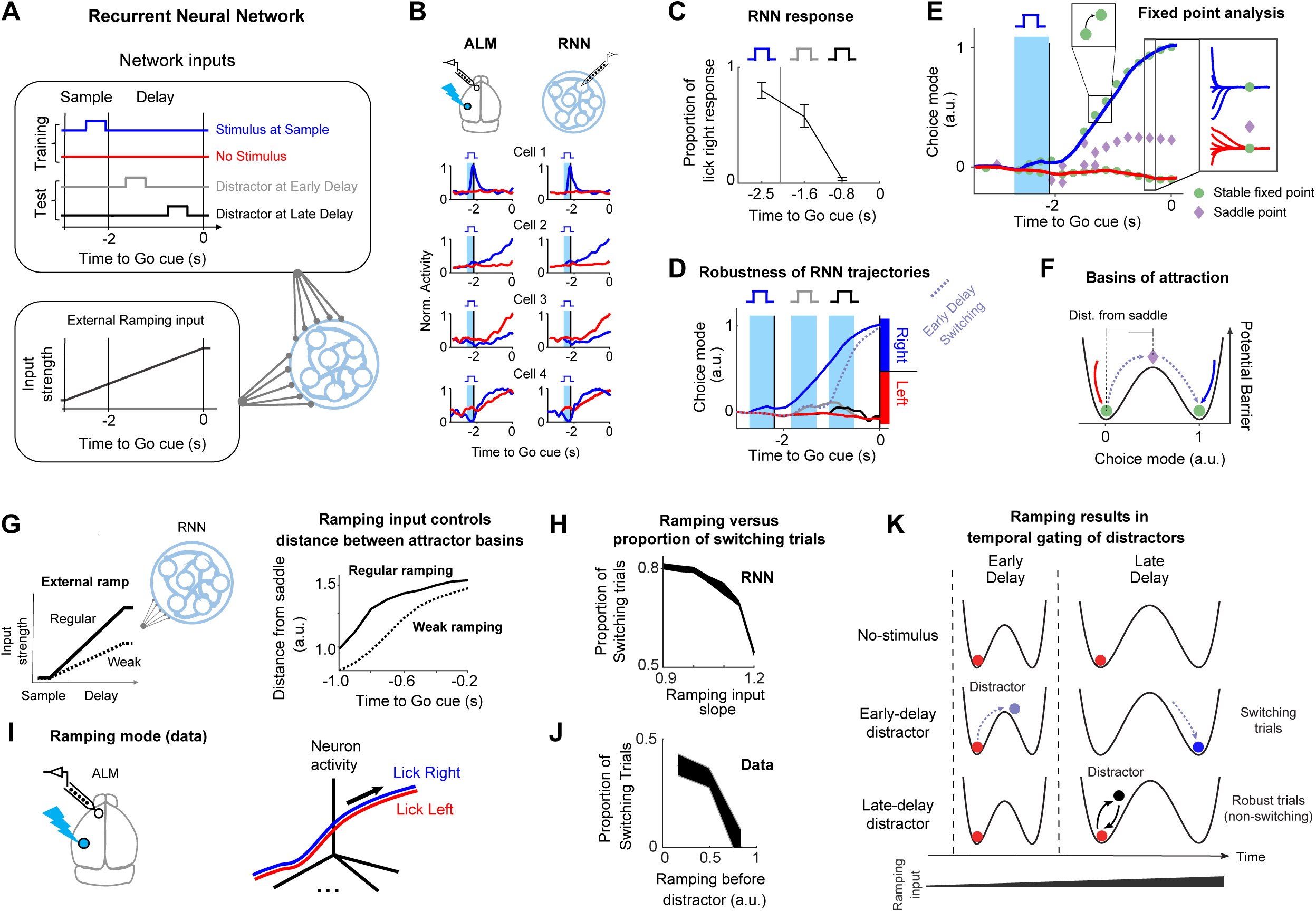
A recurrent neural network reveals temporal gating in ALM via attractor dynamics controlled by an external ramping input. **A**, ALM activity was modeled as a network of recurrently connected nonlinear rate units (RNN). A pulse delivered to a subset of units mimicked the optogenetic stimulus instructing right trials. The sample epoch was delimited by two additional inputs representing the auditory cues. A linear ramping input (black) was delivered to the network during sample and delay epochs. **B**, Each unit (right, example RNN units) was successfully trained to reproduce the PSTH of a recorded neuron (left, example ALM neurons) during correct right (blue) and left (red) trials. **C**, Proportion of right outputs generated by RNN model in response to stimulus (blue), and distractors during early (gray) and late (black) delay. The network outcome (left/right) was defined by whether the activity along the Choice mode (computed in activity space of RNN units) crossed the choice boundary at delay end (Methods). Error bars, mean ± s.e.m. across trials. **D**, RNN activity projected on Choice mode for correct trials (solid lines), color scheme as in Fig. 2E-F. Dashed-gray, trials with early-delay distractors that resulted in right response (‘switching trials’). **E**, Results of fixed-point search at all tested time points along the Choice mode: ramping input resulted in a gradual increase of the distance between fixed points during the delay. Green circles, stable fixed points. Purple diamond, saddle points. Right inset: Example trajectories starting from different initial conditions are attracted towards the fixed points. Top inset: Fixed points moved in state space as the external ramping input increased in time. **F,** Schematic of RNN phase space along Choice mode. Trajectories converge towards one of the fixed points when inside the attraction basins (solid arrows). Trajectories crossing the saddle point will end in the opposite attraction basin (dashed arrows). **G**, Left: Schematics of external ramping inputs of two different slopes (*regular* and *weak*) delivered to the RNN. Right: Distance between left fixed point and saddle point during delay, for different ramping inputs. Stronger ramping input results in larger distance to the saddle point during delay. **H**, Proportion of RNN right-output (switching trials) following early-delay distractor, as a function of ramping-input slope. Stronger ramping input reduced distractor propensity to induce a switch in RNN output. **I**, Schematics of non-selective Ramping mode in ALM. **J**, Single-trial analysis of proportion of switching trials as a function of non-selective Ramping mode amplitude in left ALM (ramping was computed before early-delay distractor, Methods). Distractors were less-capable to induce a behavioral switch (lick-right response) on trials with stronger ramping before distractors. **K**, Gating mechanism in ALM as revealed by RNN analysis. External ramping input increased the separation between attractor basins as time progressed during delay epoch. Early-delay distractor trajectories easily crossed the saddle point, resulting in frequent switches to the right basin (middle). Late-delay distractor trajectories did not reach the saddle point, and thus converged back to left basin (bottom), resulting in distractor-gating.

Selectivity in vS1 and ALM could encode sensory evidence or the chosen action (*27, 28, 8*). The same sensory input can result in different actions on correct versus error trials (**Sup. Fig. 1B**, clusters 1-3). We used neural activity on correct and error trials to decode ‘sensory’ (photostimulus) information or the upcoming ‘choice’ (lick-direction). Sensory-related information was transient and peaked first in vS1, followed by ALM (**Fig. 1H**, top). Choice-related information was nearly absent in vS1, but emerged in ALM early during delay and increased with time (**Fig. 1H**, bottom). This demonstrates that brief photostimulation caused transient sensory-related activity both in sensory and frontal cortex, whereas the frontal cortex encoded the chosen action starting in the early delay epoch.

We next asked how sensory inputs are processed as time unfolds during motor planning. We first trained mice on the basic task (as in **Fig. 1B**), and then probed mouse behavior in response to weak distractors, which consisted of weak photostimuli delivered at different times during lick-left trials (**Fig. 2A** left, Methods). These ‘distractor-naive’ mice were not exposed to distractors during training. Distractors during sample or early delay significantly increased lick-right responses compared to no-stimulus trials. In contrast, late-delay distractors did not affect behavior – demonstrating temporal gating (**Fig. 2A** right).

A different group of mice was exposed to distractors of variable strength already during training phase (‘distractor-trained’, Methods), without reward for licking right in response to distractors. This resulted in lower sensitivity to weak distractors during delay, compared to ‘distractor-naive’ mice (**Fig. 2A** right). We next probed distractor-trained mice with distractors during delay that were as strong as the sample stimulus (**Fig. 2B** left). We found that strong early-delay distractors often resulted in a right lick, while late-delay distractors did not affect behavior (**Fig. 2B** right). In other words, the same stimulus that was consistently detected during the sample epoch was ignored when delivered late in the delay epoch – demonstrating temporal gating even for strong distractors. To explain this temporal gating we considered two alternative mechanisms (**Fig. 2C**). **Hypothesis 1**: During the delay, stimulus-evoked responses become progressively attenuated in vS1 or ALM. **Hypothesis 2**: Stimuli can reach ALM and perturb the motor plan throughout the delay, but the motor plan becomes progressively more robust as the moment to act approaches. Therefore, the ability of distractors to switch the motor plan diminishes in time.

To analyze gating mechanisms on the neural level, we used dimensionality reduction (**Fig. 2D, Sup. Fig. 4** and **Methods)** to separate stimulus-related activity (‘Stimulus mode’) from activity predicting the upcoming movement (‘Choice mode’). We focused on distractor-trained mice because they exhibited prominent temporal gating (**Fig. 2B**). We first tested whether temporal gating operates early during sensory processing, by analyzing neural responses in vS1. Stimulus-evoked responses in vS1 were not attenuated during the delay epoch (**Sup. Fig. 4**, top), thus excluding gating at the level of sensory cortex. Furthermore, stimulus-evoked responses were not attenuated in ALM (**Fig. 2E, Sup. Fig. 4**, bottom), ruling out Hypothesis 1. Similar to distractor-trained mice, distractor-naive mice gated distractors in a time-dependent manner without distractor-suppression during delay epoch in vS1 or ALM (**Sup. Fig. 5**).

Analysis of the Choice mode showed that distractors delivered during delay did not initiate lick-right trajectories in ALM on trials in which the animal ignored the distractor and licked left (**Fig. 2F,** gray and black; **Sup. Fig. 6**). Instead, on trials with early-delay distractors resulting in right licks (‘switching trials’), neural activity exhibited switches from lick-left to lick-right trajectories (**Fig. 2F,** dashed gray), consistent with Hypothesis 2. Taken together, these results suggest that sensory inputs do reach ALM during the delay epoch, but the selected motor plan becomes increasingly robust to sensory input (Hypothesis 2), producing temporal gating.

To understand what circuit dynamics support temporal gating, we first trained recurrent neural networks (RNNs) to reproduce neural activity (**Fig. 3A-B**, Rajan et al., 2016). This approach preserves the rich heterogeneous dynamics observed in ALM recordings (**Sup. Fig. 1, 7**). Analysis of ALM activity showed that a significant proportion of the variance during the delay (**Sup. Fig. 3C-D)** was explained by a non-selective ramping component of the neural activity (‘Ramping mode’, Methods), which likely reflects an external input (*11, 14*). We therefore trained the network with an external ramping input (**Fig. 3A**; for analysis of a network without external ramping see **Sup. Fig. 8**). Trained RNN units produced output matching the peristimulus time histogram (PSTH) of recorded ALM neurons (**Fig. 3B, Sup. Fig. 7D-E**), during correct left or right trials without distractors. Even though the network was not trained to reproduce neural activity in trials with distractors, its dynamics showed remarkable robustness to late-delay distractors (**Fig. 3C-D**) – resembling the gating effect observed in ALM (**Fig. 2B,F**).

We reverse-engineered the RNN model (*30*) to uncover the mechanism underlying temporal gating (for analysis of the model without external ramp see **Sup. Fig. 9A-C**). During the delay epoch, small perturbations of the network state in the vicinity of left and right trajectories decayed to the original trajectories (**Fig. 3E** right inset, green circles). These dynamics indicate the presence of two point-attractors, with attractor basins separated by a saddle point (**Fig. 3E-F**, purple diamond). As the ramping input increased during the delay, it moved the attractors further apart from each other (**Fig. 3E**) and increased the distance between the lick-left trajectory and the saddle point (**Fig. 3G** right; regular ramping). Consequently, distractor stimuli became progressively less likely to overcome the distance between left and right basins of attraction (**Fig. 3K**) – resulting in temporal gating.

The separation between attractors in the RNN was controlled by the amplitude of the ramping input. When the RNN received a weaker ramping input instead of the regular ramping input (**Fig. 3G** left), the separation between lick-left attractor and saddle during delay was smaller (**Fig. 3G** right). This predicts that distractors are more likely to cause a switch on trials with weaker ramping, compared to trials with stronger ramping (**Fig. 3H, Sup. Fig. 10A-B**). Analysis of single-trial activity in ALM along the non-selective Ramping mode (**Fig. 3I**) confirmed this prediction. We grouped trials based on the non-selective ramping amplitude before distractor onset. Trials with weaker ramping before distractor onset were less robust to distractors compared to trials with stronger ramping (**Fig. 3J**; *P* = 0.003), indicating that choice robustness is controlled by ramping amplitude rather than by elapsed time *per se* (**Sup. Fig. 10A-C**).

We observed that in distractor-naive mice, which were less robust to weak distractors compared to distractor-trained mice (**Fig. 2A** right), the contribution of the non-selective Ramping mode was also smaller (**Sup. Fig. 3C-D**. For weak ramping input, the RNN model predicted that distractors can more easily push the activity beyond a saddle point, resulting in a higher proportion of switches from one attractor to the other (**Fig. 3F** dotted arrows). Indeed, strong late-delay distractors, which were ignored in distractor-trained mice, were able to induce a significant number of switches in distractor-naive mice (**Sup. Fig. 11D**). Furthermore, the impact of weak distractors on Choice mode in distractor-naive mice was more prominent and persisted for longer time than in distractor-trained mice (**Sup. Fig. 11A-C**). These results suggest that attractor basins in distractor-naive mice are shallower compared to those in distractor-trained mice (**Sup. Fig. 11E**). Taken together, our results indicate that sensory inputs are gated by attractor dynamics in ALM in a manner that is shaped by experience.

## Discussion

Decisions can flexibly change between initial action-selection and motor response (*31*–*34, 10*). We found that stimuli arriving during this gap became less capable of changing decisions with time – demonstrating a form of temporal gating. Other studies of temporal gating (*16*) suggested a mechanism based on distractor-suppression in prefrontal cortex (*35*). Our results suggest that temporal gating does not require distractor suppression, but can instead arise from a gradual increase in robustness of the chosen motor plan. This novel gating principle complements previously observed mechanisms by which temporal expectations modulate attention during stimulus selection (*36*–*40*).

To infer the mechanism underlying temporal gating, we trained recurrent neural networks (*41, 18, 39, 42, 43*) to reproduce the heterogeneous dynamics of individual neurons (*29*) that we recorded in ALM. We found that RNN models were able to generalize to conditions they were not trained to reproduce, and predicted temporal gating as observed in the data. Reverse-engineering the trained RNN models (*30*) revealed a novel mechanism for temporal gating, which operated via a combination of attractor dynamics and a ramping signal.

Attractor dynamics can underlie short-term memory (*44*–*54*), preparatory activity (*11, 14*), and potentially support distractor-resistant memory (*55*–*57, 11, 14*). Here we propose that attractor dynamics in ALM provide an effective gating mechanism that regulates motor-plan flexibility in response to incoming inputs. This mechanism can explain previous studies showing that relevant sensory inputs, distractors, and direct perturbations of decision-making circuits have stronger influence on behavior when applied early during the trial (*16, 58, 35, 52, 40, 59*). Because neural activity in ALM can be changed by experience, the dynamical structure that emerged in this task could be different in other decision-making paradigms (*60*–*63, 18, 64*–*66, 42, 67, 68*).

Our work also suggests a new computational role for the ramping signal, which has been observed in many decision-making tasks and is related to time estimation and a sense of urgency (*69, 70, 7, 5, 71, 72, 14*). Ramping might act as a temporal discounting signal for incoming sensory inputs by gradually increasing the commitment to the selected action (*34*), by moving attractors apart over time. Taken together, our work uncovered a general mechanism for temporal gating that allows for robust neural representation of the motor plan until the moment to act.

## Acknowledgments

We thank David Hansel and Shaul Druckmann for discussions; Johnatan Aljadeff, Kayvon Daie, Ran Darshan, Jan Drugowitsch, Hidehiko Inagaki, Marton Rozsa, and Tim Wang for comments on the manuscript; Tina Pluntke for animal training.

## Authors Contributions

AF, MNE, and KS designed the experiments, with help from NL. AF and MNE collected the experimental data with help from NL. AF analyzed the behavioral and neural data, with input from MNE, LF, SR, and KS. LF and SR conceived the computational models with input from AF and KS. LF performed network simulations. AF, LF, SR and KS wrote the paper, with input from all the authors.

## Data and materials availability

Data and code for simulations will be made available.

## Methods

### Animals

We used 9 adult Scnn1a-TG3-Cre × Ai32 transgenic mice, which express Channelrhodopsin-2 (ChR2H134R-EYFP) in layer 4 stellate cells in the barrel cortex (Ai32 mice, JAX 012569; Scnn1a-TG3-Cre mice, JAX, 009613) (*25, 73*–*75*). All experimental procedures were approved by Janelia Institutional Animal Care and Use Committee. Detailed procedures for water restriction and surgical procedures were described elsewhere (*8, 14*).

### Mouse behavior

Water-restricted mice were trained to report detection of photostimulation by directional licking for a water reward.

#### Basic task (Fig. 1B)

Mice were head-fixed in front of two lick ports and were instructed to lick right or left for a water reward. Each trial began with a pre-sample epoch (1.2 s duration), followed by a sample epoch (1 s duration) that was demarcated by auditory cues (0.15 s duration, ‘sample cues’). Photostimulation during the sample epoch instructed mice to lick-right. Absence of photostimulation during the sample epoch instructed mice to lick-left. Mice were trained to withhold licking for an additional 2 s after the sample epoch (delay epoch), until an auditory ‘Go cue’ (pure tone, 3.4 kHz, 0.1 s duration) signaled the beginning of the response epoch (1.5 s duration). Early licks (occurring before the Go cue), were punished by restarting the delay epoch.

#### Task with distractors (Fig. 2A-B, left)

Distractors were photostimuli outside of the sample epoch. Mice were not rewarded for licking right in response to distractors. Lick-left trials with distractors resulting in right licks (which were considered error trials) were referred to as *switching trials* (for example, **Fig. 2F**, dashed gray ‘switching trials’). We used two groups of mice: ‘distractor-naive’ (*n* = 5) and ‘distractor-trained’ mice (*n* = 4). Initially all mice were trained on the basic task without distractors until reaching criterion (75% correct trials). ‘Distractor-trained’ mice were then trained to explicitly ignore distractors, ‘distractor-naive’ mice were trained only on the basic task. During training of distractor-trained mice the amplitude of the distractor photostimuli was gradually increased until it was identical to that of the stimulus. In half of the trials, distractors were also delivered on lick-right trials (data not shown). In each session ∼75% of the trials included distractors. During recording sessions, both distractor-naive and distractor-trained mice were tested in the presence of distractors of different amplitudes and timings (see next section ‘Photostimulation’ for details).

### Photostimulation

Photostimuli from a 473-nm laser (Laser Quantum) were controlled by an acousto-optical modulator (Quanta Tech). Photostimuli were delivered to vS1 (anteroposterior (AP) −1.3 mm; medial lateral (ML) 3.5 mm, relative to bregma) on the left hemisphere, through a clear skull cup (*8*). A guide tube (1.6 mm inner diameter) was glued on top of the stimulation site, and an optical fiber was inserted into the guide tube at the beginning of each behavioral session. During sessions in which we combined electrophysiological recordings from vS1 with photostimulation, the laser beam was positioned directly above the recording site. To prevent mice from utilizing visual cues to distinguish photostimulation trials from control trials, a masking flash (10 Hz) was delivered using 470-nm LEDs throughout the trial.

We used 3 types of photostimuli (**Fig. 2A-B**):

1. ‘Stimulus’ was delivered during the sample epoch (at −2.5 s before the Go cue) and instructed mice to lick right. It contained 4 pulses lasting 0.4 s (10 Hz, sinusoidal temporal profile). ‘Stimulus’ peak power was typically 2.2 mW, but in a few sessions power was adjusted to correct for performance biases (mean power = 2.2 mW, std = 0.2 mW, range [1.2-3.2] mW, across sessions).
2. ‘Strong Distractors’ had the same photostimulation profile as the ‘Stimulus’, but were delivered outside of the sample epoch (onsets = −3.5, −1.6, −0.8 s relative to Go cue).
3. ‘Weak distractors’ contained one pulse that lasted for 0.1 s, with peak power typically set to be 33.3% of the ‘Stimulus’ peak power (mean power = 0.8 mW, std = 0.2 mW, range [0.4-1.1] mW, across sessions). ‘Weak-distractor’ photostimulus power was calibrated prior to the first recording session, to ensure that it elicited a change in behavioral-performance when delivered during the sample epoch. Weak distractors were delivered during the sample or delay epoch (onsets = −3.5, −2.5, −1.6, −0.8 s relative to Go cue).

### Electrophysiology

We recorded extracellular spikes using silicon probes (2 shanks × 32 channels) with 250 µm spacing between shanks and 25 µm spacing between channels (H2; Cambridge Neurotech). The 64-channel voltage signals were multiplexed, recorded on a PCI6133 board (National Instrument), and digitized at 14 bits. The signals were demultiplexed into 64 voltage traces, sampled at 25 kHz and stored for offline analyses. Recordings were done through a small craniotomy (diameter, 0.5 −1.0 mm) made one day before the recording. The craniotomy was centered over left or right ALM (anteroposterior (AP) 2.5 mm; medial lateral (ML) 1.5 mm, relative to bregma) or left vS1 (−1.3 mm AP; 3.5 mm ML). The brain was allowed to settle for at least 10 min following penetration with the silicon probe, before the recording started. Depths of recorded neurons in ALM (ranging between 250 and 1,350 µm) and vS1 (between 175 and 1,150 µm) were inferred from manipulator readings. We performed 3-10 recording sessions from each craniotomy on consecutive days. Distractor-naive mice: left ALM (*n* = 26 sessions), right ALM (*n* = 11), and vS1 (*n* = 10). Distractor-trained mice: left ALM (*n* = 23), right ALM (*n* = 11), and vS1 (*n* = 16).

Behavioral and electrophysiological data was stored and analyzed using custom pipeline in DataJoint framework (*76*).

### Behavioral data analysis

For behavioral analyses we excluded early-lick trials (trials during which the animal started licking before the Go cue) and no-response trials (trials that did not elicit any licking until 1.5 s after the Go cue). These trials were also excluded from analysis of neural activity. Behavioral performance for each session was calculated as the proportion of lick-right responses on each trial type, and expressed as mean ± s.e.m. across sessions. Data in **Fig. 1C** is shown for distractor-naive mice. For distractor-trained mice the proportion of lick-right responses was: 0.79 ± 0.02 on stimulus (lick-right) trials, 0.25 ± 0.01 % on no-stimulus (lick-left) trials (*n* = 4 mice, 87 sessions).

To assess the effects of stimulus/distractors on performance (**Fig. 2A-B** right) we calculated, for each session, the change in proportion of lick-right responses on trials with stimulus or distractors relative to control trials (i.e. trials with instruction to lick left, without distractors **Fig. 2A-B** left, red):

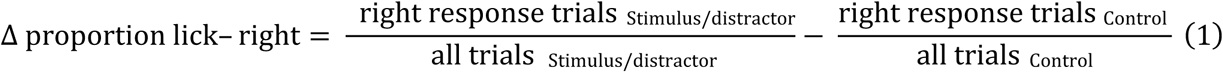

The control trials accounted for the proportion of spontaneously occurring lick-right trials in the absence of stimulus or distractors. Data in **Fig. 2A-B** is based on: distractor-naive mice with weak distractors (*n* = 5 mice, 67 sessions); distractor-trained mice with weak distractors (*n* = 4 mice, 19 sessions); distractor-trained mice with strong distractors (*n* = 4 mice, 67 sessions).

### Electrophysiological data analysis

Spike sorting was done using JRClust (*77*). A total of 3385 well-isolated single units were recorded of which we included 3011 based on the following criteria: (i) The cell had a mean spike rate equal to or larger than 0.5 Hz. (ii) The cell was recorded for at least 10 lick-left and 10 lick-right correct trials without distractors. In addition, we recorded 1799 multi-units, of which we included 1611 units based on the criteria above.

Putative pyramidal neurons and fast-spiking interneurons were differentiated based on spike-waveforms (*8, 14*). Putative pyramidal cells recorded in: left ALM (1648 well-isolated neurons; 769 multi units), right ALM (679; 365), and left vS1 (256; 291). Putative fast-spiking interneurons recorded in: left ALM (184 well-isolated neurons; 21 multi units), right ALM (96; 13), and left vS1 (92; 116).

All analyses were performed on putative pyramidal neurons, except for the analyses of putative fast-spiking interneurons presented in **Sup. Fig. 2**. For all population analyses we combined single units with multi-units, except for the analyses presented in **Sup. Fig. 1-2**. We only analyzed trials with ≥ 5 simultaneously recorded units (number of units per trial = 45.3 ± 0.1, mean ± s.e.m. across trials). All analyses in **Fig. 1** (except panel **C**) and **Sup. Fig. 1-2** were performed on data from distractor-naive and distractor-trained mice combined.

### Population-average spike rate and selectivity

*Spike rates* were computed in 5 ms time bins, with spike-time in each trial defined relative to the Go cue. Trial-average spike rates were smoothed with a 50 ms causal boxcar filter. For plotting PSTH of example cells and population averages we used a 50 ms (**Fig. 1D-E, Sup. Fig. 2A, Sup. Fig. 4A, Sup. Fig. 5B**), or 200 ms filter (**Sup. Fig. 6A, Sup. Fig. 11A**).

*Grand-average population responses* (**Fig. 1F-G, Sup. Fig. 2A**) were computed by averaging the trial-average spike rates across cells, using correct lick-left or lick-right trials without distractors.

Trial typ*e selectivity S*_*i*_(*t*) at time *t* of cell *i* was defined as:

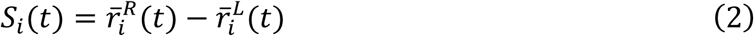

where 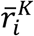 are the trial-average spike rates at time t, computed using correct right (*K* = *R*) or left trials (*K* = *L*).

Neurons were identified as trial type selective during the sample or delay epochs if their epoch-averaged selectivity was significantly different from zero (Mann–Whitney U test, *P* ≤ 0.001). To determine selectivity of neuron *i* during sample epoch we averaged *S*_*i*_(*t*) in a 0.5 s long window following the stimulus onset. To test for significant selectivity during delay epoch we averaged *S*_*i*_(*t*) during the entire 2 s of the delay.

### Hierarchical clustering of neural activity profiles in response to photostimulation

We used hierarchical clustering (implemented using MATLAB functions *pdist, linkage*, and *cluster*, where the distance between observations was based on pairwise correlations) to categorize trial-averaged neural activity responses across recorded cells (**Sup. Fig. 1A-B, Sup. Fig. 2B-C**). For each cell we concatenated the trial-average spike rates for both lick-left and lick-right trial types, computed using correct trials. Trial-average spike rates were smoothed with a 200 ms causal boxcar filter. Clustering was done based on correlation between trial-average responses across cells. For clustering we used a time window of t = [-3.5, 0] s, which included sample and delay epochs. Putative pyramidal and fast-spiking interneurons were clustered separately. In ALM, clustering was done using well-isolated cells recorded from left and right ALM. In vS1 we combined well-isolated cells and multi-units together.

For display purposes (**Sup. Fig. 1A-B, Sup. Fig. 2B-C**), we averaged the spike rate across all neurons that belonged to the same cluster, separately for correct and error trials. For correct trials we used only trials without distractors. For error trials (which were less frequent) we combined error trials from all trial types, including trials with distractors. Only cells with at least 5 trials of each trial type were included. Only clusters with more than 2 % of cells are shown.

### Decoding of sensory- and choice-related neural activity

To decode sensory- and choice-related neural activity (**Fig. 1H**) we trained a support vector machine classifier (implemented using MATLAB function *fitcsvm* with linear kernel function). To decode sensory-related activity we labeled the trials based on the instruction (i.e. whether a stimulus was present during sample epoch). To decode choice-related activity, we labeled the trials based on whether the mouse licked left or right, irrespective of the instruction. We trained the classifier on matched number of correct and error trials in both lick directions, to ensure that the instruction is decorrelated from the outcome. Specifically, each training set was composed of 4 categories of trials (correct left, error left, correct right, and error right), with the same number of trials in each category. Matching was performed by subsampling: we randomly selected the same number of trials from all categories (subsampling was repeated 20 times), with the number of trials selected being equal to the size of the smallest category. We included sessions that i) had at least 5 simultaneously recorded ‘stable’ cells, including multi-units, ii) with at least 10 trials in the smallest category, including trials with distractors. In each session, for each subsample, we trained the classifier based on 80 % of the selected correct and error trials, and tested it on the remaining 20 % of trials. This process was repeated 100 times. The decoder performance was taken as the average decoder performance across all repetitions. Decoding was done in 0.1 s time bins, with the classifier being retrained for each time bin. For decoding we used only ‘stable’ cells – i.e. cells that were recorded on at least 80% of the trials within each session.

### Population dynamics in neural activity space

We analyzed the population dynamics of *n* simultaneously neurons recorded in a session. **(Fig. 2E-F, Sup. Fig. 3, Sup. Fig. 4D, Sup. Fig. 5C** bottom, **Sup. Fig 6B, Sup. Fig. 11B)**. During each trial, the population activity of *n* neurons drew a trajectory in the *n*-dimensional activity space, where each dimension represents the spike rate of one neuron. We identified directions (*modes*) in the activity space that maximally separated the neural trajectories for different trial conditions and times during the trial (**Sup. Fig. 3A**)(*11, 14*). When analyzing responses to distractors in the activity space, we restricted the analysis to trial types that had at least 5 neurons recorded simultaneously with at least 5 trials per condition.

*Stimulus mode* was computed as a *n* × 1 vector of trial-averaged spike rate differences of *n* neurons during trials with lick-right and lick-left *instructions*, averaged within a 0.5 s window following the stimulus onset during sample epoch:

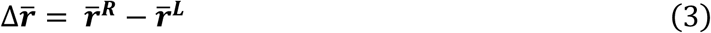

The resulting vector (Eq. 3) was normalized by its *ℓ*_2_ norm, resulting in a vector of weights (one weight per neuron):

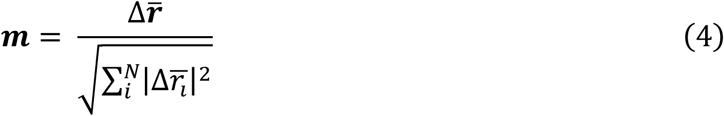

Projections of the neural activity along the mode over time were calculated as:

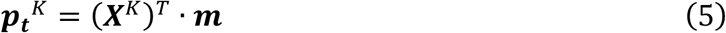

Where ***X***^*K*^ is *n x t* matrix of spike rates of neurons over time, for trial type *K*. Normalizing each mode by its *ℓ*_2_ norm (Eq. 4) ensured that projected activity ***p***_***t***_^*K*^ would not scale with the number of neurons recorded simultaneously. If an individual neuron was not recorded on a particular trial, its weight in (Eq. 4-5) was set to zero when computing the projected neural activity for that trial.

*Choice mode* was defined as a *n* × 1 vector of trial-averaged spike rate differences of *n* neurons during trials with lick-right and lick-left *outcomes*, averaged within a 0.2 s window at the end of the Delay epoch, before the Go cue (analogously to Eq. 3-4). The number of correct and error trials in both lick directions was matched to ensure that the instruction is decorrelated from the outcome. Specifically, we used 4 categories of trials (correct left and error right trials – both resulting in lick-left outcome; correct right and error left trials – both resulting in lick-right outcome), with the same number of trials in each category. Matching was performed by subsampling: we randomly selected the same number of trials from all categories (subsampling was repeated 20 times), with the number of trials selected being equal to the size of the smallest category. If a session did not have at least 10 trials in the smallest category (including trials with distractors), we matched the number of trials with left and right outcome, irrespective if they came from correct or error trials. To compute the Choice mode using left-preferring cells (**Sup. Fig. 6B**, right) we set the positive weights of the Choice mode to 0, and then projected the neural activity on this mode as described above. Positive weights corresponded to right preferring cells, whereas negative weights corresponded to left-preferring cells (Eq. 3, 4).

We also identified a non-selective *Ramping mode* (**Fig. 3I, Sup. Fig. 3A-B**), which captured the difference in neural activity (regardless of the trial type) between the beginning of the trial and the end of delay epoch. We computed the Ramping mode similarly to the Choice and Stimulus mode: we took the difference in trial-averaged spike rate (but pooling together both lick-left and lick-right trials) during the last 0.5 s of the Delay epoch versus the 0.5 s that preceded the sample epoch:

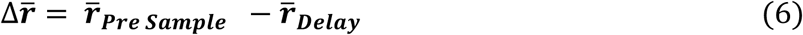

The modes were orthogonalized to each other using a Gram-Schmidt process in the following order: Choice, Ramping, Stimulus mode.

To compare trial-averaged projections across sessions, we first subtracted from each trial-averaged projection the baseline activity computed in the same session. Baseline activity was defined as the trial-averaged activity computed on correct lick-left trials without photostimulation, averaged over the 1 s window preceding the sample epoch. Trial-average projections were smoothed with a 100 ms causal boxcar filter for display. After averaging the projected activity across sessions, the resulting averaged projection was normalized to its maximum value reached at any time before the Go cue on lick-right correct trials. Projections along the Stimulus and Ramping modes were computed by combining correct and error trials, with at least 15 trials per trial type. Projections along the Choice mode were computed separately for correct trials and error trials, with at least 5 trials per trial type and outcome.

### Variance explained by different modes

To calculate the fraction of variance explained by neural activity projected on each mode for each session (**Sup. Fig. 3C-D**), we first performed baseline subtraction from the spike rate of each neuron. Specifically, we subtracted the trial-averaged baseline spike rate of each neuron from the spike rate on each trial. Baseline spike rate of each neuron was computed by averaging the spike rate across all trials over 1 s window that preceded the sample epoch. We used the same baseline-subtracted spike rates to project the neural activity on the different modes. The projections to each mode were smoothed with a causal sliding boxcar window of length 0.5 s with 0.1 s steps. For Stimulus and Ramping modes we used correct and error trials without distractors. For Choice mode we used only correct trials, without distractors.

To calculate the *fraction of trial-to-trial variance explained* (**Sup. Fig. 3C**) by each mode, we calculated the trial-to-trial variance in each time bin as a sum (across neurons and trials) of square of spike rate averaged in each time bin. We divided this value by the trial-to-trial variance of the neural activity projected on a given mode, calculated as the sum (across trials) of squares of the projection averaged in each time bin:

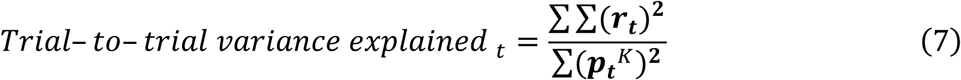

To calculate the *fraction of trial-averaged variance explained* (**Sup. Fig. 3D**), we calculated the total trial-averaged variance in each time bin as a sum (across neurons) of squares of the trial-average spike rate left and right trial types, averaged in each time bin. We divided this value by the square of the trial-average projection in left and right trial types, averaged in each time bin:

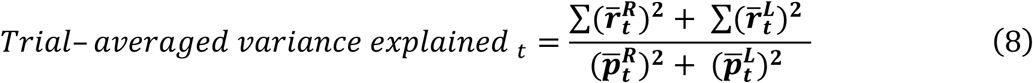

### Distractor response-size quantification

We computed the changes in spike rate of neurons in response to photostimulation (*Δ spike rate*, **Sup. Fig. 4B, Sup. Fig. 5C** top) as follows: For each session, we first computed the *grand population averaged spike rates* (see definition in section Population-average spike rate and Selectivity), separately for each trial type, with correct and error trials pooled together. To get the *Δ spike rate* in each time bin, we took the difference between the *grand population averaged spike rate* for each trial type, and that on lick-left trials without distractors.

The *Response size* was defined as the number of spikes added during photostimulation (*Δ spikes*, **Sup. Fig. 4C, Sup. Fig. 5D** top). For each session we averaged *Δ spike rate* during the time window of the photostimulation (strong distractors, 0.4 s time window; weak distractors 0.1 s time window).

Because the number of responsive cells was smaller in ALM compared to vS1, we also quantified the change in ALM activity during photostimulation using the Stimulus mode (*Stimulus mode response*, **Sup. Fig. 4E, Sup. Fig. 5D** bottom) – which allowed to detect changes in neural activity during photostimulation. For each session and for each trial type with photostimulation, we projected trial-averaged neural activity on the Stimulus mode, combining correct and error trials. From this, we subtracted the trial-averaged projection on lick-left trials without distractors. The resulting change in projection activity was averaged over a given time window during photostimulation (stimulus and strong distractors, 0.4 s time window; weak distractors 0.1 s time window). The *Stimulus mode response* to distractors at different times/strength was normalized to the *Stimulus mode response* to the sample (i.e. to photostimulation presented during sample epoch).

We compared photostimulation response size in vS1 and ALM at different time-points (**Sup. Fig. 4C,E**, **Sup. Fig. 5D**) using repeated measures analysis of variance (ANOVA) and found no significant differences in responses to photostimulation at different time points during the task (*P* > 0.05).

### Distractor impact on Choice mode

To assess the impact of distractors on the Choice mode (**Sup. Fig. 6C, Sup. Fig. 11C**), for each session we computed trial-averaged projections of neural activity on the Choice mode on trial types with distractors. We aligned the trajectories to distractor onset and subtracted the projection on control correct trials (lick-left trials without distractors). This was done separately for trials with distractors delivered during early-delay or late-delay. We then averaged the activity across trial-averaged projections corresponding to different distractor times. We limited this analysis to 0.8 s from distractor onset – the minimal common duration for each distractor trajectory until the Go cue (the time interval from the late-delay distractor onset and the Go cue). The resulting trajectory was defined as the *distractor impact* on Choice mode (**Sup. Fig. 6C, Sup. Fig. 11C)**.

### Single-trial analyses of the Choice mode

To analyze the separation of neural activity projected on the Choice mode (**Sup. Fig. 6D-E**) on lick-left versus lick-right trials, we analyzed the distribution of projections on single trials, averaged over the last 0.2 s of the delay epoch (*end-delay points*). To compare across sessions, the distribution of end-delay points for each session was normalized (*14*): the median of end-delay points on lick-left trials was set to 0, and the median on lick-right trials was scaled to 1. To compute the medians, we used all response trials (including distractors), with lick-left and lick-right trials referring to the trial outcome. **Sup. Fig. 6D** shows end-delay points distribution (± s.e.m. across sessions) on correct lick-left versus lick-right trials without distractors. In **Sup. Fig. 6E** (right) we computed the end-delay points distribution (± s.e.m. across sessions) for trials with early-delay distractors, in which the animal ignored the distractor and licked left (‘robust trials’, gray) and trials in which the animal licked-right (‘switching trials’, light blue). Analogously, in **Sup. Fig. 6E** (left), we computed the distribution of trajectories on robust and switching trials, during mid-delay ([−0.8, −0.6] s, relative to Go cue). This time window corresponded to the middle of the interval between the onset of the early-delay distractor (−1.6 s, relative to Go cue) and the Go cue.

*Decoding of behavioral outcome from the neural activity projected along the Choice mode (***Sup. Fig. 6F)**. For each session, we first computed the optimal classification threshold for single-trial projections of lick-right versus lick-left trials via the Receiver Operating Characteristic method (ROC). We averaged the projections over the last 0.2 of the delay epoch. Lick left or lick right class labels were assigned to each trial according to the corresponding behavioral outcome (including distractor trials). Then, for each session, we counted the proportion of projected trials that could be classified as resulting in a lick right response using the optimal threshold, after subtracting the proportion of no-stimulus trials that spontaneously resulted in a right lick (as we did for behavioral analyses in **Fig. 2B**).

### Single-trial analyses of the Ramping mode

To analyze if ramping predicted the degree of robustness to distractors (**Fig. 3J**) we analyzed population activity projected on the non-selective Ramping mode (Eq. 6, **Sup. Fig. 3A-B**) on single trials. For each trial, we smoothed the projected neural activity with a 200 ms causal boxcar filter. We limited our analysis to early-delay distractor trials, for which there were many switching trials. In each session, for a given trial, we computed the *ramping amplitude* before an early-delay distractor as the ramping-mode projection averaged in a 0.4 s time window ending before the onset of the distractor (−1.6 s). We binned the distribution of *ramping amplitudes* across trials into 3 equally sized bins, corresponding to weak, intermediate, and strong ramping amplitudes. For each bin, we computed the proportion of distractor trials with a lick-right outcome. We performed analogous computations for different ramping levels on control trials (i.e. trials with instruction to lick left, without distractors) – to obtain the proportion of spontaneously occurring lick-right trials. To compute the ramping values on control trials, we used the same time window that we used for computing ramping values before distractor on distractor trials. For each session we subtracted the proportion of lick-right responses on control (no distractor) trials from the proportion of lick-right responses on distractor-trials, computed for corresponding ramping bins. The resulting difference was defined as proportion of switching trials for each level of non-selective ramping (x).

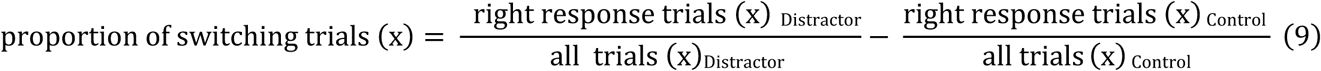

Subtracting the control distribution allowed to estimate the proportion of switches caused by distractors, by accounting for the proportion of spontaneous lick-right responses that occurred in trials without distractors. Statistical significance of the effect of ramping amplitude on the proportion of switching trials was assessed using repeated measures analysis of variance, ANOVA [F (2, 24) = 7.2, *P* = 0.003].

### Choice-mode trajectory slope on correct and switching trials

To compute the Choice mode projection slope (**Sup. Fig. 8G**) on correct lick-right trials versus switching early-delay distractor trials, we computed the trial-average projection of neural activity along the Choice mode for each of the trial types. We then aligned the trajectory to stimulus/distractor onset (**Sup. Fig. 8D-F**). Slope was computed as the difference in trial-averaged trajectory averaged over a time window t_1_ = [-0.2, 0] s and time window t_2_ = [1.4, 1.6] s, divided by the difference in time window midpoints (t_2_ – t_1_ = 1.6 s). Time was defined relative to stimulus/distractor onset. These time windows were chosen, because for early-delay distractors, they spanned the entire period between distractor onset and the end of delay epoch. The same calculation was carried on to determine the slope of the trajectories generated by the RNN (see below).

### Recurrent neural network model

To study ALM circuits while maintaining the heterogeneity of individual neuron responses during trials, we built a rate-based recurrent neural network in which the output of each unit matched the PSTH of an experimentally recorded unit in left ALM of distractor-trained mice pooled across sessions. We trained two classes of RNNs: i) an autonomous RNN where the slow ramping dynamics observed in the data must be generated via recurrent dynamics, and ii) a RNN receiving an additional, linearly ramping, external input which can be integrated by the network to generate the slow timescale seen in the data.

### Network dynamics, inputs to the network and connectivity

The networks were composed of *N* rate units whose membrane currents, ***x***, were modeled with the following system of *N* coupled first-order differential equations:

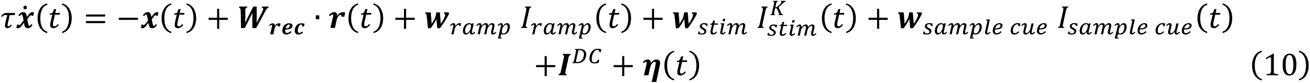

The spike rate was computed from the membrane currents by applying a sigmoidal transduction function elementwise:

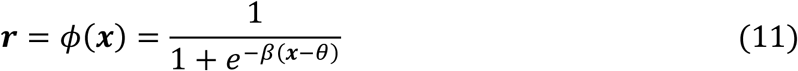

The parameters of *ϕ*(***x***) were set to *β* = 0.8 and *θ* = 3, to generate a Gaussian distribution of currents from the skewed lognormal distribution of spike rates (**Sup. Fig. 6A**). The decay time constant *τ* was 10 ms for all units. ***η***(*t*) was a low-pass filtered white noise:

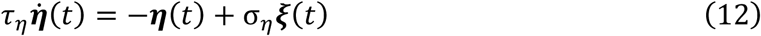

with *τ*_*η*_ = 2 ms. White noise ***ξ***(*t*) was drawn at each time step with standard deviation 1. *σ*_*η*_ was 0.15 during both training and testing of the RNN models. Higher values of *σ*_*η*_ (> 0.5) produced a single stable fixed point, instead of bistable dynamics (data not shown). All numerical solutions of the network dynamics were obtained using the first order Euler-Maruyama method with time step of Δ*t* = 0.1 ms. Synaptic weights from a given unit in the network could be both excitatory and inhibitory. The initial value of the recurrent synaptic matrix was:

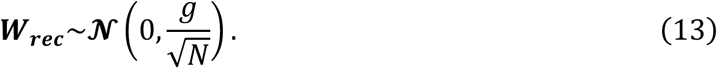

Since all synaptic weights were trained, the value of *g* did not influence the dynamics at the end of the training procedure (we trained networks with *g* = [0.9 − 1.5] and did not observe substantial qualitative differences in the network dynamics post-training). Constant task-independent inputs were drawn from a Gaussian distribution of mean 0 and standard deviation 1, ***I***^*DC*^∼**𝒩**(0,1). We used three weight vectors for the different task-dependent inputs supplied to the network: ***w***_*stim*_ for stimulus and distractors, ***w***_*ramp*_ for external ramping, ***w***_*sample cue*_ for the external cues delimiting the sample epoch. The weight vectors were drawn from three Gaussian distributions: ***w***_*stim*_∼**𝒩**(0,1), ***w***_*ramp*_∼**𝒩**(0,1), ***w***_*sample cue*_∼**𝒩**(0,0.1). 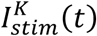 was zero during lick-left trials (*K* = *L*) and had a square pulse profile (duration *T* = 400 *ms*) during lick-right trials (*K* = *R*). The stimulus started at 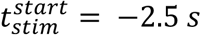 for the regular stimulus delivered during right trials, at 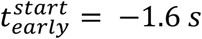 for early-delay distractor trials, 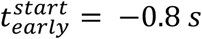 for late-delay distractor trials. *I*_*sample cue*_(*t*) had a square pulse profile (duration *T* = 150 *ms*). The first auditory cue was delivered at 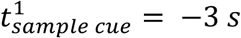 before the Go cue, the second at 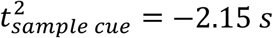. In the RNN of **Fig. 3** the external input *I*_*ramp*_(*t*) was a linearly ramping function of time, which started to ramp at the beginning of the sample epoch 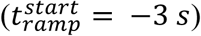 until the Go cue (*t*_*go cue*_ = 0 *s*). *I*_*ramp*_(*t*) was set to zero for the RNN with autonomous dynamics (**Sup. Fig. 8-9**). Auditory cues, and the nonspecific external input were identical during lick-right and lick-left trials. At each trial, all time-dependent inputs were subject to a random jitter drawn from a uniform distribution within the interval [-10,10] ms to mimic variability of information transfer from other brain areas to ALM.

### Network training procedure

The networks learned to reproduce the activity of single recorded units at each timepoint throughout the trial (*29*). We defined the total input of each unit in the network at time *t* as:

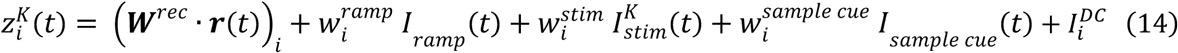

At each time step and for each unit the error is computed as the difference between the current input of a unit and its target function: 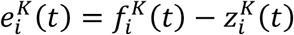. To build the individual target function 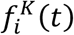 of unit *i* we: i) averaged the spike rate of a recorded neuron at time *t*, across lick-right (*K* = *R*) or lick-left correct trials (*K* = *L*); ii) normalized the average spike rate with the function:

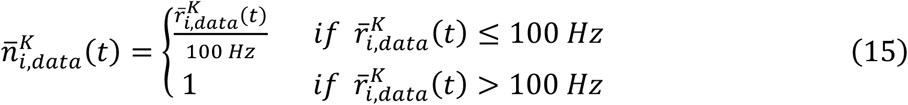

iii) obtained the membrane current by inverting Eq. 11: 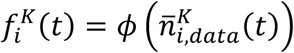. To build the target functions and to run RNN simulations we only considered the last 500 ms of the pre-sample, the sample and the delay epochs, excluding the response epoch. We did not consider data recorded outside of these epochs. All time-dependent inputs and target functions were further smoothed with a 400 ms boxcar moving window. After smoothing, units that displayed an average spike rate lower than 0.01 Hz in at least one timebin (resolution of 1ms) were not included in the RNN models. The total number of units, including both single- and multi-units, amounted to *N* = 668. We only considered units from distractor-trained mice recorded in left ALM.

Incoming synapses to unit *i* were changed according to the first-order reduced and controlled error (FORCE, Sussilo & Abbott 2009) algorithm:

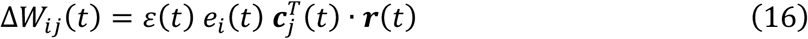

where ***c***_*j*_(*t*) is the *j*th column of ***C***(*t*), a running estimate of the inverse of the correlation matrix of network rates, plus a regularization term: ***C***(*t*) = (∑_*t*_ ***r***(*t*) · ***r***(*t*)^T^ + *α****I***)^−1^. The prefactor *ε*(*t*) plays the role of an effective learning rate at each time step, and is defined as:

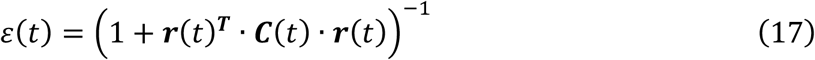

The initial state ***C***(0) was set to *α* × ***I*** with *α* = 1 (Sussillo and Abbott, 2009), ***I*** is the identity matrix. The slope of the ramping input current *I*_*ramp*_(*t*) during training was drawn from the uniform distribution with support [*μ*_*ramp*_ − 0.1, *μ*_*ramp*_ + 0.1] with *μ*_*ramp*_ = 1. The peak amplitude of the stimulus current 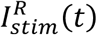 was drawn at each trial from the uniform distribution in the range [*μ*_*stim*_ − 0.1, *μ*_*stim*_ + 0.1] with *μ*_*stim*_ = 1. The synaptic update of Eq. 16 was not performed at every timestep to avoid overfitting. Instead, in each trial we sampled intervals between training time bins from a random uniform distribution with support [70 − 35, 70 + 35]*ms*, which was sufficient to avoid overfitting single-trial fast noise. The learning algorithm was stopped after 500 trials, when the input to each unit closely matched their target functions (**Sup. Fig. 7E**). At the beginning of a given trial, a trial type label (lick-right or lick-left) was randomly assigned to that trial (with its corresponding input *I*_*stim*_(*t*)).

### Simulation of ALM dynamics with trained recurrent neural networks

We first generated 100 lick-right (stimulus during sample epoch, **Fig. 3A** blue input) and 100 lick-left (no stimulus, **Fig. 3A** red input) trials from the RNN to obtain the Choice mode. In each trial both the initial conditions, the jitters and the fast noise realizations were randomized. The Choice mode was computed as the vector of spike rate differences during lick-right and lick-left trials averaged during the last 400 ms of the Delay epoch 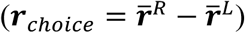. To avoid biases in the estimate of the Choice mode, we excluded aberrant trials (<10% of all trials), i.e. trials that exhibited fluctuations six or more times bigger than the standard deviation of lick-right or lick-left trials at any timepoint during the Delay epoch. In these simulations the amplitude of the fast noise *σ*_*η*_ was set to zero and the the stimulus amplitude ramping input was set to the same value used during training. The ramping slope in the RNN with external ramping input was drawn from a gaussian distribution of mean *μ*_*ramp*_ = 1 and standard deviation *σ*_*ramp*_ = 0.1. After estimation of the Choice mode, we further simulated 200 lick-right and 200 lick-left trials. Trial *k* was considered to be correct if its trajectory, projected onto the Choice mode 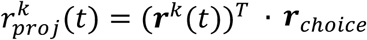, lied above the halfway point between 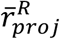 and 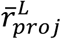 at the end of the Delay epoch (3.5s into the trial corresponding to the time of the go cue, **Fig. 3D**). A trial was defined as an error trial if its projected trajectory would lie below the halfway point.

To investigate whether the network was able to generalize to conditions that were not presented during training we probed it with early and late distractors (200 trials each, **Fig. 3C**). Trials were classified as correct or switching using the same criterion described above for establishing the Choice mode. A small (<10%) fraction of trials displayed large fluctuations and diverged significantly from either lick-right or lick-left typical trajectories; these aberrant trials were not included in our analyses. In these simulations the stimulus/distractor amplitude was drawn from a gaussian distribution with mean *μ*_*stim*_ = 1 and standard deviation *σ*_*stim*_ = 0.4; the slope of the external ramping input was drawn from a gaussian distribution of mean *μ*_*ramp*_ = 1 and standard deviation *σ*_*ramp*_ = 0.1.

### Fixed point analysis

To discover the mechanism generating the network dynamics we ‘reverse-engineered’ the RNN (Sussillo & Barak, 2013). We examined the noiseless dynamics of the network at time 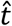, which is governed by the right-hand side of Eq. 10:

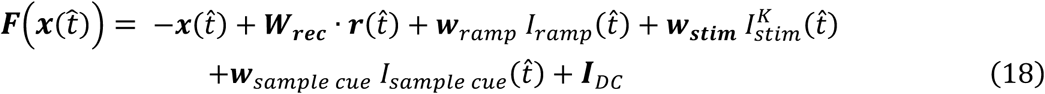

We looked for points in the activity space spanned by all network units where the dynamics was slow by first expanding 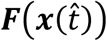 around candidate points:

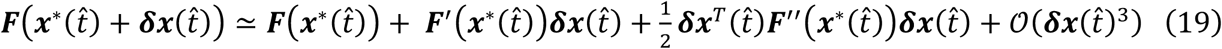

Our analysis encompassed fixed points, i.e. point where 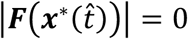, and slow points, i.e. points where 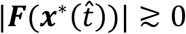, but the linear term 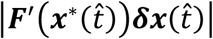 dominated over all other terms in the Taylor expansion of Eq. 19. This can be implemented by minimizing the scalar convex function (Sussillo & Barak, 2013):

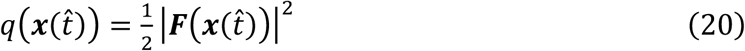

where the vertical bars indicate *ℓ*_2_ vector norm.

At a given time point, we set the state of the network at a given distance *ϵ* from the cross-trial average correct lick-left trajectory and calculated the value of 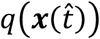, after having adjusted the external input to their mean values at time 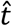 (**Sup. Fig. 9A,D**). We restricted the range of the search by shifting the network state only along the Stimulus mode or the Choice mode (note, however, that the fixed point would not necessarily be found on either of these modes). The tolerance level for fixed points (**Fig. 3E**) was set to *q*(***x***^*^) = 10^−10^. To assess the stability of such points we examined the linearized dynamics around them, and deemed the point *stable* if all eigenvalues of the Jacobian matrix 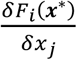 were negative semidefinite, or deemed the point a *saddle* if any of the eigenvalues were found to be positive. For slow points (**Sup. Fig. 9B**) the tolerance level was set to *q*(***x***^*^) = 10^−1^. In their vicinity, the network dynamic would evolve on a time scale significantly slower than the single neuron decay time (*τ* = 10 *ms*), confirming the presence of a slow manifold marked by those points.

### Dynamics of autonomous RNN and of RNN with external ramping input

Fixed point analysis of the autonomous RNN revealed the presence of two stable fixed points near the end-delay points of correct lick-right and lick-left trajectories. In this network the two fixed points were separated by slow points (**Sup. Fig. 9B**); while this mechanism explained the slowly ramping activity observed in the network, it could not reproduce switching trajectories as observed experimentally (**Sup. Fig. 8D,F**). The slope of switching trials (**Sup. Fig. 8G**) was calculated using the same procedure applied to the experimental data (see section ‘Choice-mode trajectory slope on correct and switching trials’ above).

The presence of the additional external input allowed RNN trained with an external ramping input to find a different solution to the problem of generating slow ramping dynamics. Here, lick-left and lick-right correct trajectories remained close to a pair of stable fixed points during the entire Delay epoch and were separated by a single saddle point (**Fig. 3E**). As time unfolded, both the distance between the two stable fixed points and the distance between the saddle and the lick-left stable fixed points increased. Ramping input slopes, regular = 1.0, weak = 0.9. Distance to saddle point increased from t = −1.2 s, which corresponds to the time of early delay distractor offset.

The strength of the external ramping input moved the fixed points apart from each other and therefore controlled the increase in distance (**Fig. 3G**). Consequently, the proportion of switching trials provoked by early-delay distractors was inversely related to the slope of the external input: stronger ramping led to increased robustness (**Fig. 3H**). To further explore this relationship, we measured the proportion of switching trials as we systematically varied the slope of the ramping input and the time at which we delivered the distractor during the Delay epoch. Both decreasing the slope of the ramping input and moving the distractor forward in the Delay epoch caused more errors (**Sup. Fig. 10B**). This suggests that elapsed time and amplitude of ramping input are directly related: the gating of late distractors can be explained by the increased amplitude of the external stimulus compared to earlier times in the delay. We performed a control analysis where for *t* > −1.2*s* we fixed the external ramping input to its value at the offset of the early-delay distractor *I*_*ramp*_(*t* = −1.2*s*) and measured the error rate in trials with early- and late-delay distractors. We found no significant difference in the proportion of switching trials between early and late distractors when the external ramping input was constant (**Sup. Fig. 10C**). This result confirmed our hypothesis on the role of ramping as the central mechanism for temporal gating. To further characterize the attraction basins we measured the relaxation time constants of network activity along the Choice mode in the RNN with external ramping (**Sup. Fig. 10D**). We first set the ramping input to the value reached at time *t* in the delay, then we waited for the network’s activity to set on no-stimulus trajectory (corresponding to lick-left) and we delivered a perturbation of amplitude 0.2 along the Choice mode. We measured the relaxation constant *τ* by fitting a single exponential decay function 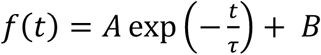 to the decaying trajectories. The relaxation time diminished as ramping slope increased, suggesting a deepening of the lick-left attraction basin with increased ramping (or, equivalently, towards the end of the delay).

### Robustness to quenched noise in recurrent weights

Both network types, with and without external ramping input, displayed some degree of robustness to multiplicative synaptic noise. We applied multiplicative gaussian noise with zero mean and *σ*_*W*_ = 0.03 to all recurrent weights. We simulated 100 realization of the network with 200 stimulus and 200 no-stimulus trials in each realization. More than 50% of the realizations showed similar behavior (proportion of correct/switching trials) and dynamics (comparable end-delay points range) to the original network, both in the external ramping and in the autonomous case. With *σ*_*W*_ = 0.05 only 20% of network realizations would exhibit dynamics compatible with ALM recordings.

**Supplementary Figure 1.**
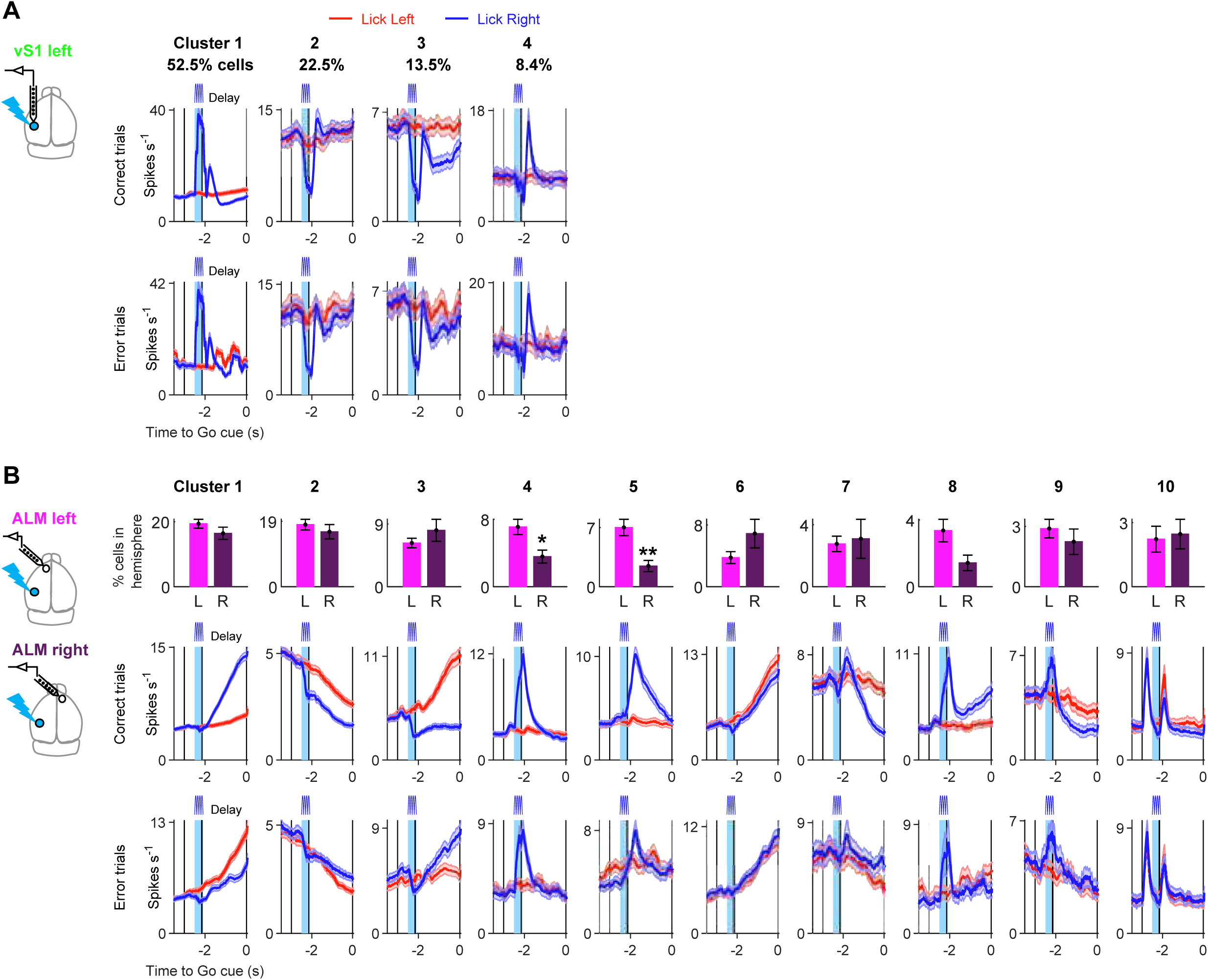
Dynamics of putative pyramidal neurons in the basic task. Hierarchical clustering of trial-averaged PSTHs of putative pyramidal cells recorded in vS1 (**A**) and ALM (**B**). **A,** Average spike rate of vS1 cells in each cluster and percentage of cells in each cluster. Red: trials without photostimulation, (lick-left). Blue: trials with photostimulation (lick-right); Cyan bar indicates the photostimulus. Top, correct trials. Bottom, error trials. Data is presented as trial-averaged activity, then averaged across cells belonging to each cluster ± s.e.m. (across cells, shaded). Cells in vS1 (clusters 1-4) did not switch selectivity on error trials, indicating that they tracked stimulus-related information. **B,** Same for ALM neurons. Neurons recorded from both hemispheres were clustered together. Bars show the percentage of putative pyramidal cells in left and right ALM that fall within each cluster. Error bars, mean ± s.e.m., across sessions; * *P* < 0.05, ** *P* < 0.01 by Mann–Whitney U test. Cells in ALM often showed opposite selectivity on error trials (e.g., Cluster 1-3), indicating evidence for preparatory activity related to choice (*13*). Despite the laterality of the movement, trial type selective preparatory activity was similar in left and right ALM (see distribution of cells in clusters 1-3 across hemispheres). Only a minority of cells in ALM did not switch selectivity on error trials – indicating sensory responses (e.g., Cluster 4; note that cells in this cluster were more prevalent in left hemisphere). In addition, there was a cluster with neurons exhibiting non-selective ramping responses (Cluster 6), and a cluster with neurons responding to auditory cues that delineated the sample epoch (Cluster 10).

**Supplementary Figure 2.**
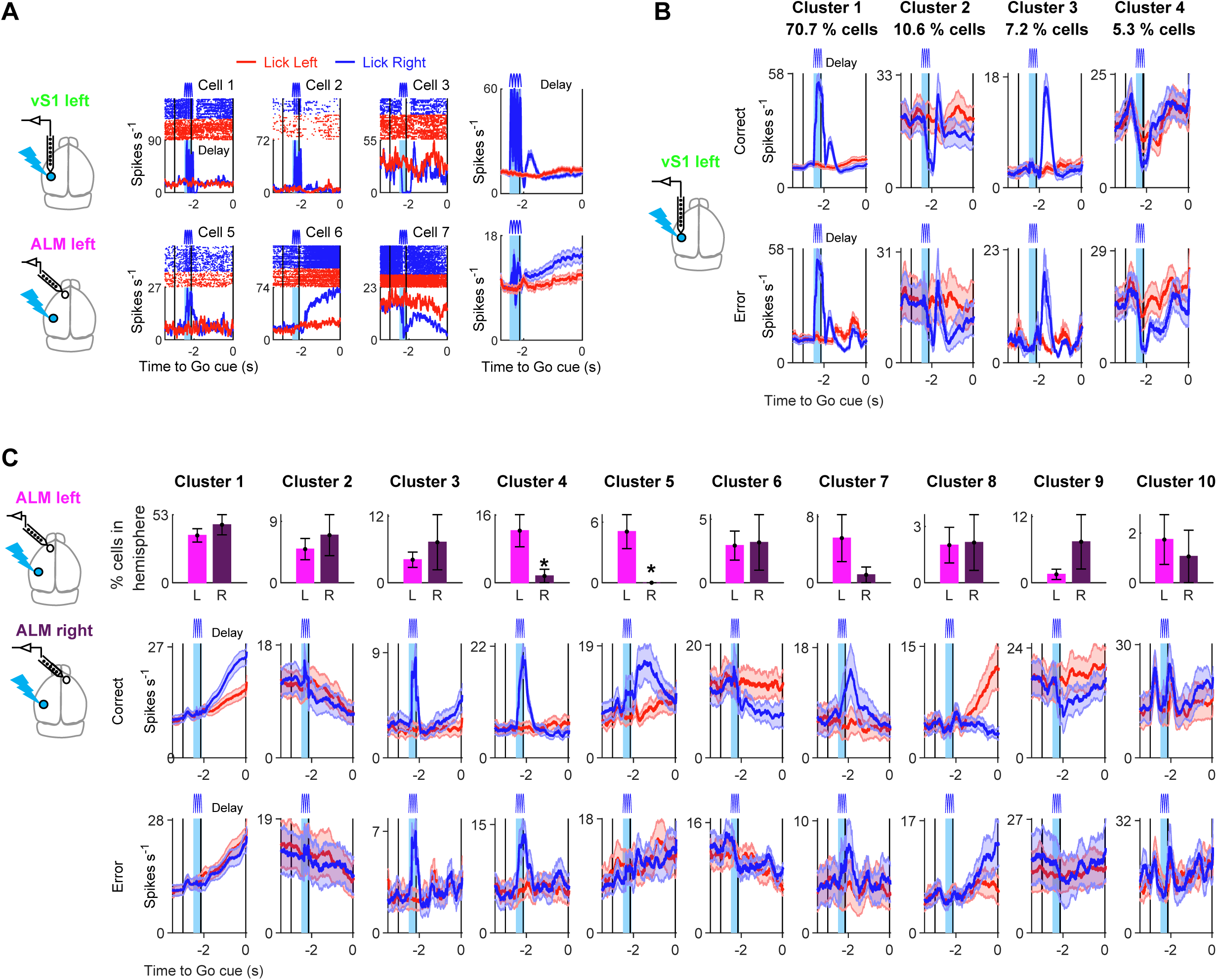
Dynamics of putative fast-spiking interneurons in the basic task. Activity of putative fast-spiking interneurons (**Methods**). **A,** Example cells and grand population average (labels as in Fig. 1D-G). **B-C,** Hierarchical clustering of trial-averaged PSTHs of putative interneurons (labels as in Sup. Fig. 1 A-B). Putative interneurons in vS1 and ALM show a diversity of response profiles, suggesting that their activity does not simply reflect average activity of excitatory neurons.

**Supplementary Figure 3.**
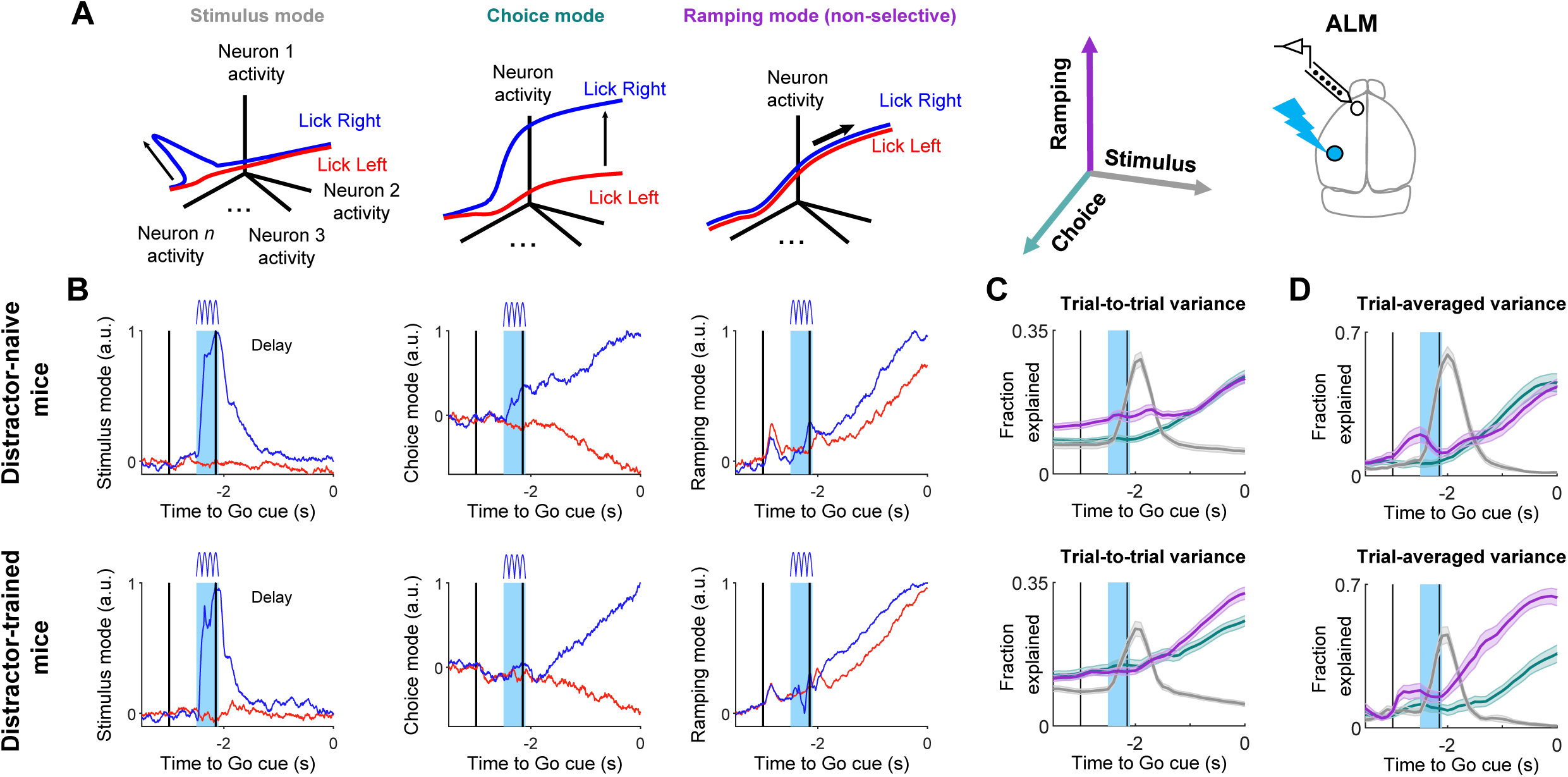
Dimensionality-reduction on population dynamics in ALM. **A,** Targeted dimensionality-reduction to define Stimulus, Choice (trial type selective), and Ramping (non-selective) modes in neural activity space. Modes are defined as one-dimensional subspaces in activity space (arrows) and are orthogonalized with respect to each other (Methods). **B**, Session-averaged projections of neural activity in left ALM on different modes in distractor-naive (top) and distractor-trained (bottom) mice. Red, lick-left trials; blue: lick-right trials. **C,** Proportion of trial-to-trial variance explained by different modes in distractor-naive (top) and distractor-trained (bottom) mice. Variance was computed at different time-points along the trial. **D,** Same as in C, for proportion of trial-averaged variance explained. Trial-averaged variance was computed separately for lick-left and lick-right trials. In all plots, for Choice mode analyses we used correct trials, excluding trials with distractors; for Stimulus and Ramping modes analyses we used correct and error trials, excluding trials with distractors.

**Supplementary Figure 4.**
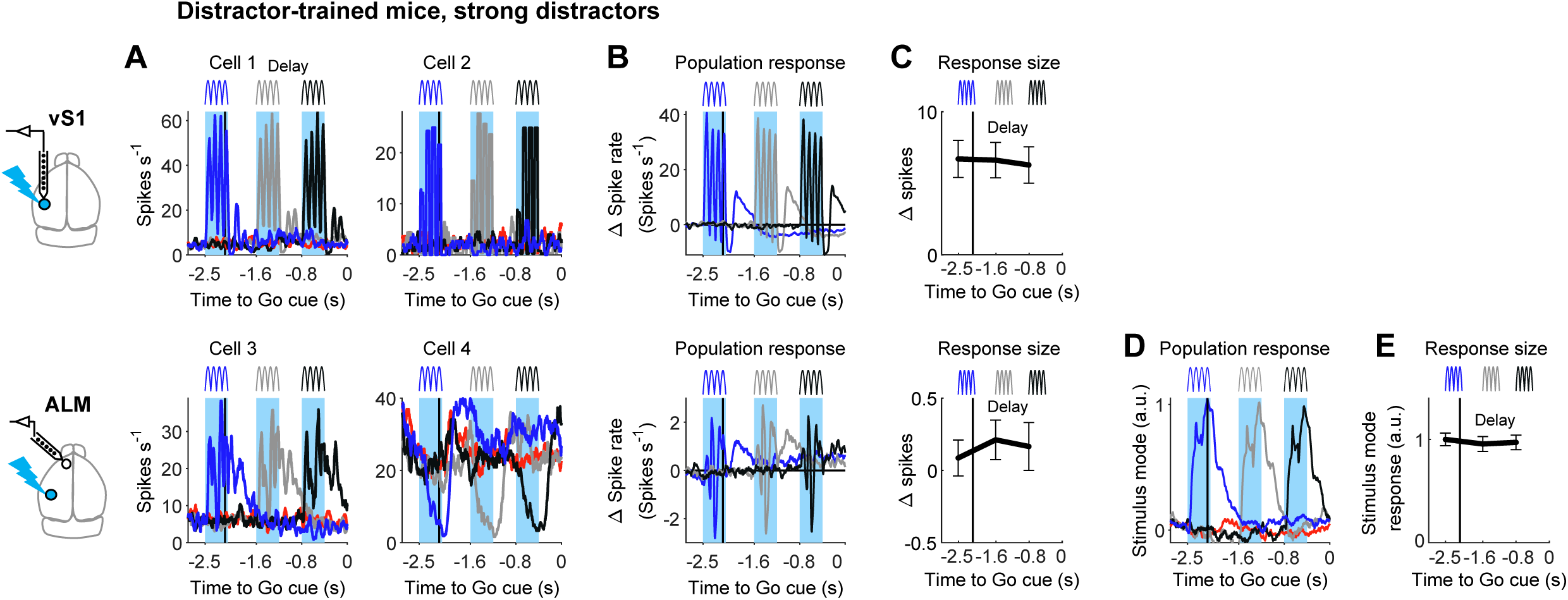
Transient activity in vS1 and ALM in response to photostimulations in distractor-trained mice. Spike rate modulation in vS1 (top) and left ALM (bottom) of distractor-trained mice in response to strong distractors at different times, computed using all response trials, regardless of the behavioral outcome. **A**, Spike rates of example cells. **B**, Population-averaged responses. **C**, Quantification of the response size based on the population average (B). Lick-left trajectory without stimulation (red), with distractors during early-delay (gray), or late-delay (black). Lick-right trajectory during sample-epoch stimulation (blue). **D-E,** For ALM recordings, we also used targeted dimensionality reduction to extract stimulus-related activity (**D**, ‘Stimulus mode’, Methods). Response to distractors projected on Stimulus mode was normalized to the response to stimulus during the sample epoch (**E, Methods**). That there was no reduction of distractor size in vS1 or ALM of distractor-trained mice (P>0.05 by repeated measures ANOVA), despite differences in the effect of distractors on behavior (Fig. 2B).

**Supplementary Figure 5.**
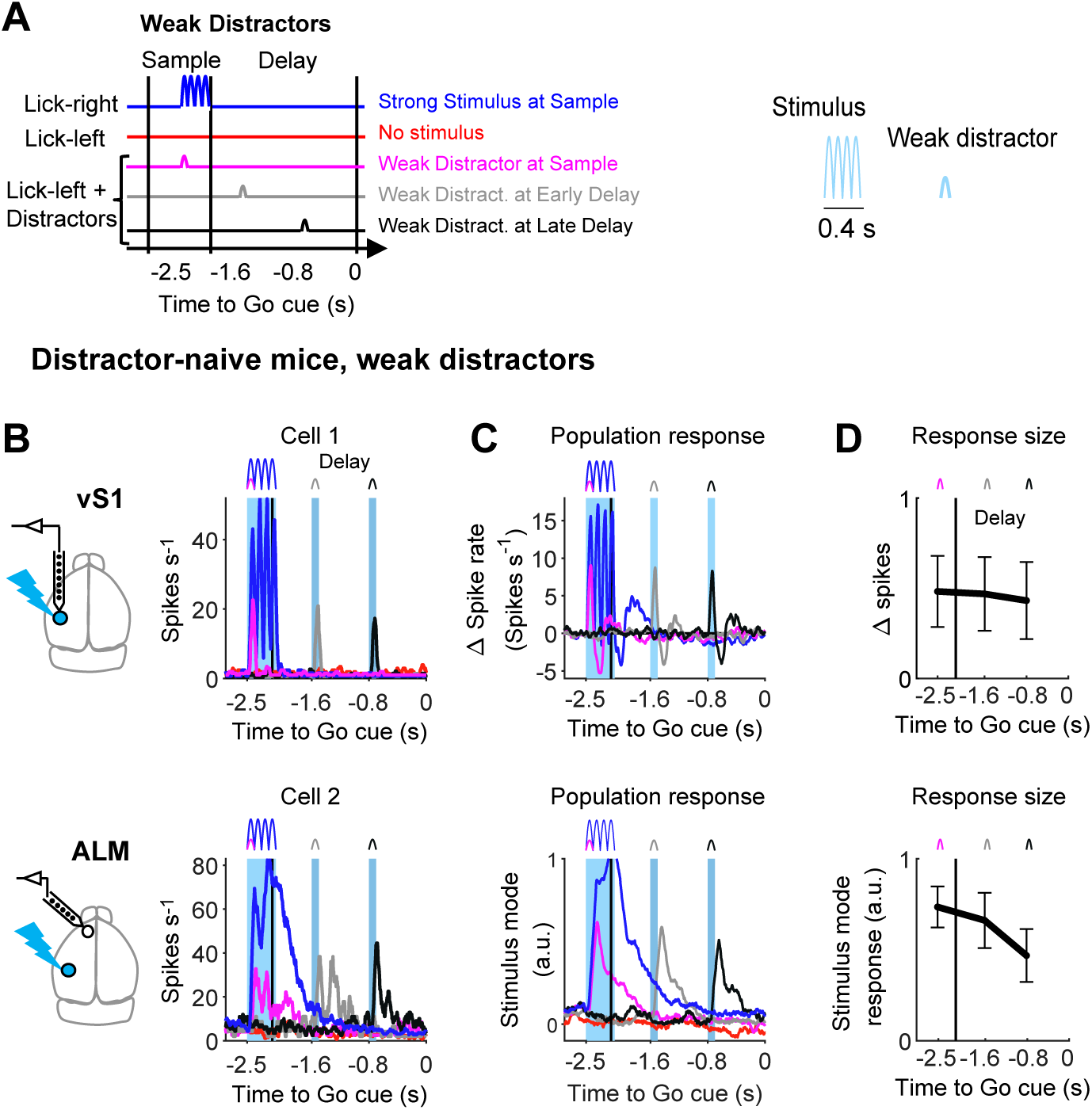
Transient activity in vS1 and ALM in response to photostimulations in distractor-naive mice. **A**, Task with weak distractors (for behavioral effect see Fig. 2A right). Transient spike rate modulation in vS1 (top) and left ALM (bottom) of distractor-naive mice in response to weak distractors at different times, computed using all response trials, regardless of the behavioral outcome. **B,** Example cells. **C**, Population responses. **D**, Quantification of response size. Response size was measured using the population average in vS1 (C-D, top), and Stimulus mode in ALM (C-D, bottom). Response to distractors projected on Stimulus mode was normalized to the response to stimulus during the sample epoch (**E, Methods**). Lick-left trajectory without stimulation (red), and with weak distractors during sample (magenta), early-delay (gray), or late-delay (black). Lick-right trajectory during sample-epoch stimulation (blue). There was no reduction of distractor size in vS1 or ALM of distractor-naive mice (P>0.05 by repeated measures ANOVA).

**Supplementary Figure 6.**
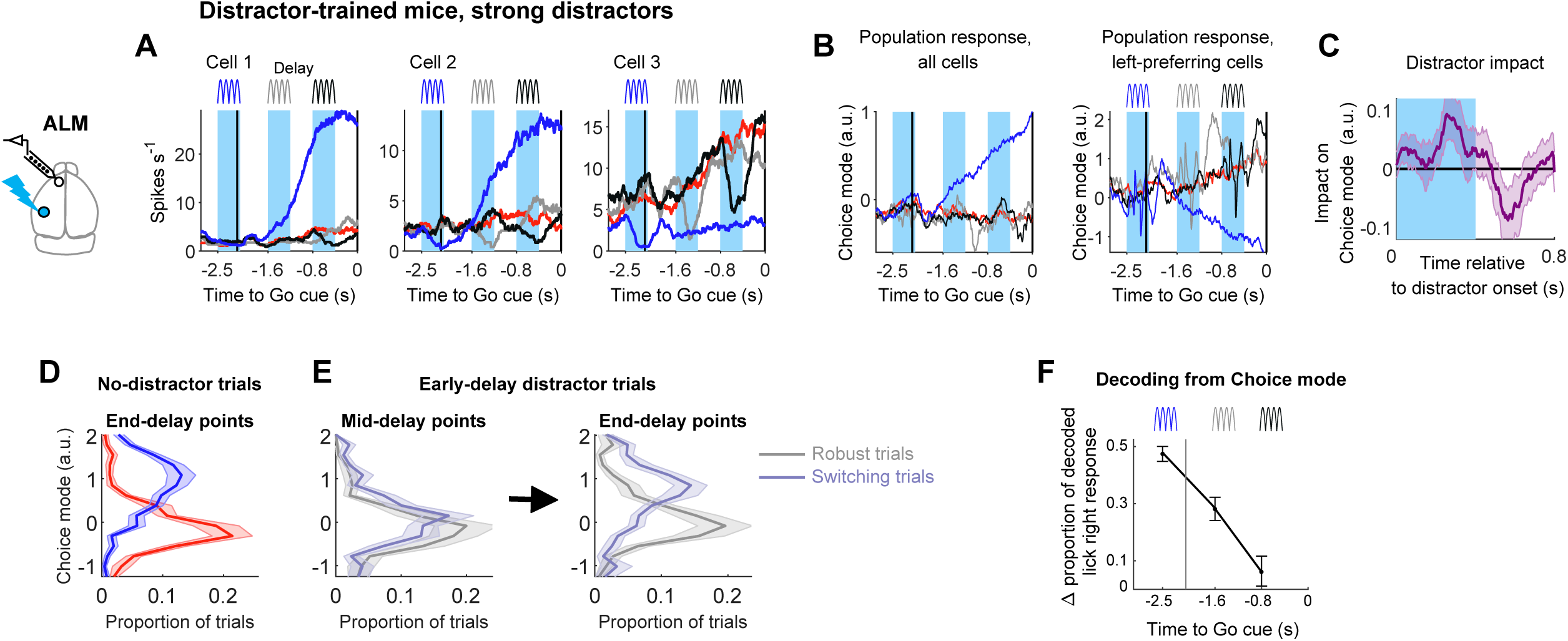
Robustness of the Choice mode in response to distractors. Spike rate modulation in ALM in response to strong distractors at different trial epochs, in distractor-trained mice. **A,** Example cells. **B**, Projection of neural activity on the Choice mode using all cells (left panel) or using only left-preferring cells (right panel). Lick-left trajectory without stimulation (red), with distractors during early-delay (gray), or late-delay (black), computed using correct trials. Lick-right trajectory during sample-epoch stimulation (blue). **C,** Impact of distractors on the Choice mode. Trajectories were aligned to the onset of each distractor. Data is shown as average across sessions and distractors with different onset times ± s.e.m. (shaded). Data in A-C was computed using correct trials. The effect of distractors on the Choice mode on correct trials was temporary: distractors often resulted in transient change in spike rate of individual cells that contributed to the Choice mode, but their activity later recovered to the unperturbed (red) trajectory, indicating robustness. **D,** Distribution of Choice mode projections on single trials without distractors, averaged over the last 0.2 s of the delay epoch (*end-delay points)* for lick-left (red) versus lick-right (blue) trials. **E,** Distribution of Choice mode projections on trials with early-delay distractor (gray, robust trials; light blue, switching trials). Projections were averaged during mid-delay (left panel) or end-delay (right panel, Methods). Data is expressed as mean ± s.e.m. (shaded) across sessions. **F,** Proportion of lick-right responses on distractor trials (relative to control trials), decoded from neural activity on single trials projected on the Choice mode. This result confirmed that neural trajectories along the Choice mode were more robust to late-delay distractors compared to early-distractor, with a high correspondence between the temporal gating of distractors observed on the neural and behavioral level (compare with **Fig. 2B**).

**Supplementary Figure 7.**
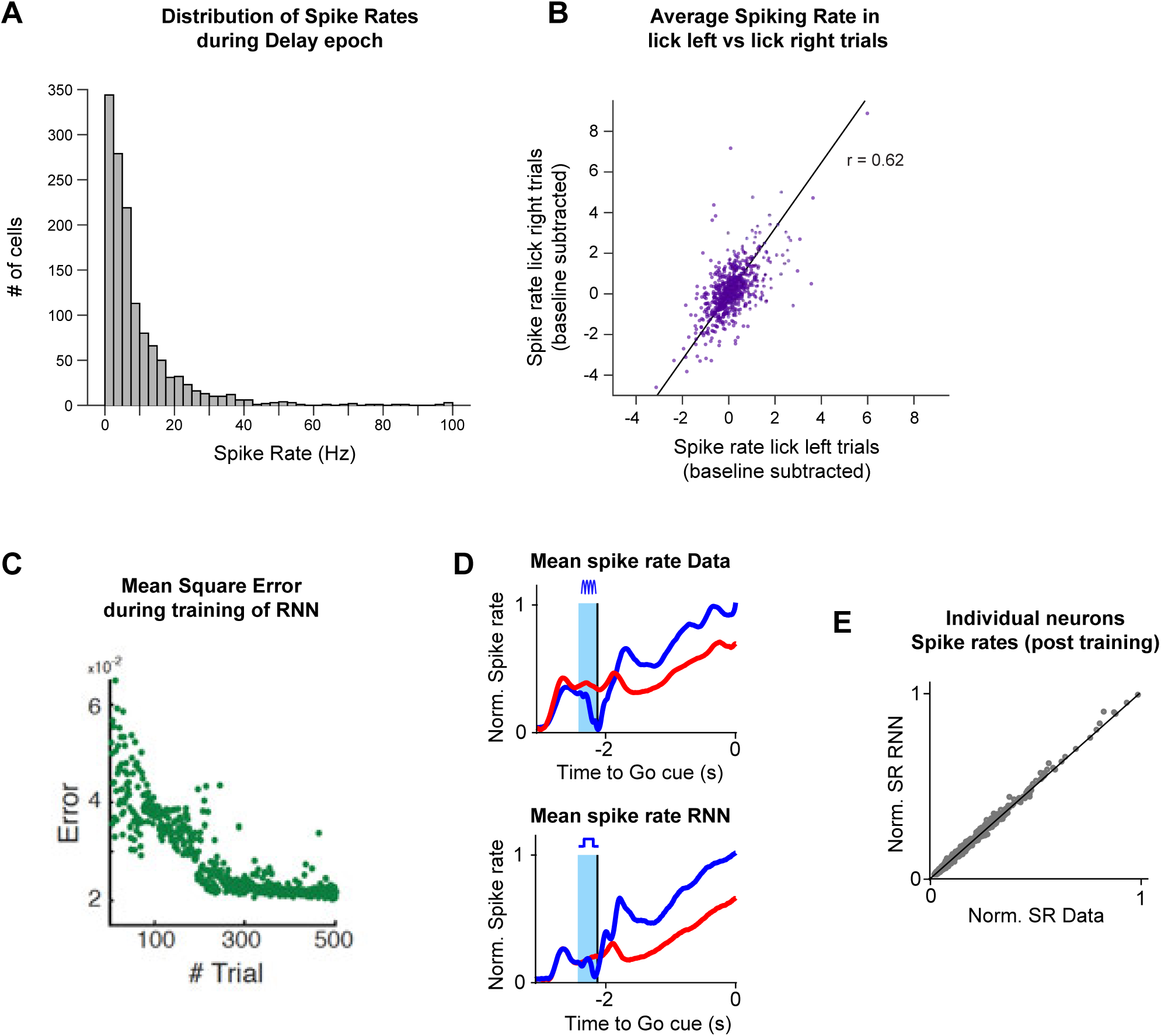
Heterogeneous neuronal spike rates in ALM and results of RNN training. **A,** Distribution of spike rates across units in ALM, averaged during the delay epoch. **B,** Trial-averaged spike rates during lick left versus lick right trials. Each point represents the spike rate of a unit during delay epoch, after subtraction of the average baseline spike rate of that unit. represents the Pearson correlation coefficient. **C-E,** Outcome of training procedure of RNN with external input. **C**, Mean square error between the normalized PSTHs of target ALM neurons and the output of corresponding RNN units during training. The training procedure was terminated after 500 trials. **D**, Spike rate of recorded ALM units (top) and RNN units at the end of training (bottom), averaged across units and trials and normalized by the maximum spike rate across time and units. **E**, Normalized spike rates of all units incorporated in the RNN, averaged during the delay epoch, data versus RNN (post-training). All points lie near the diagonal, indicating that at the end of the training procedure RNN units correctly reproduce the activity of neurons recorded in ALM.

**Supplementary Figure 8.**
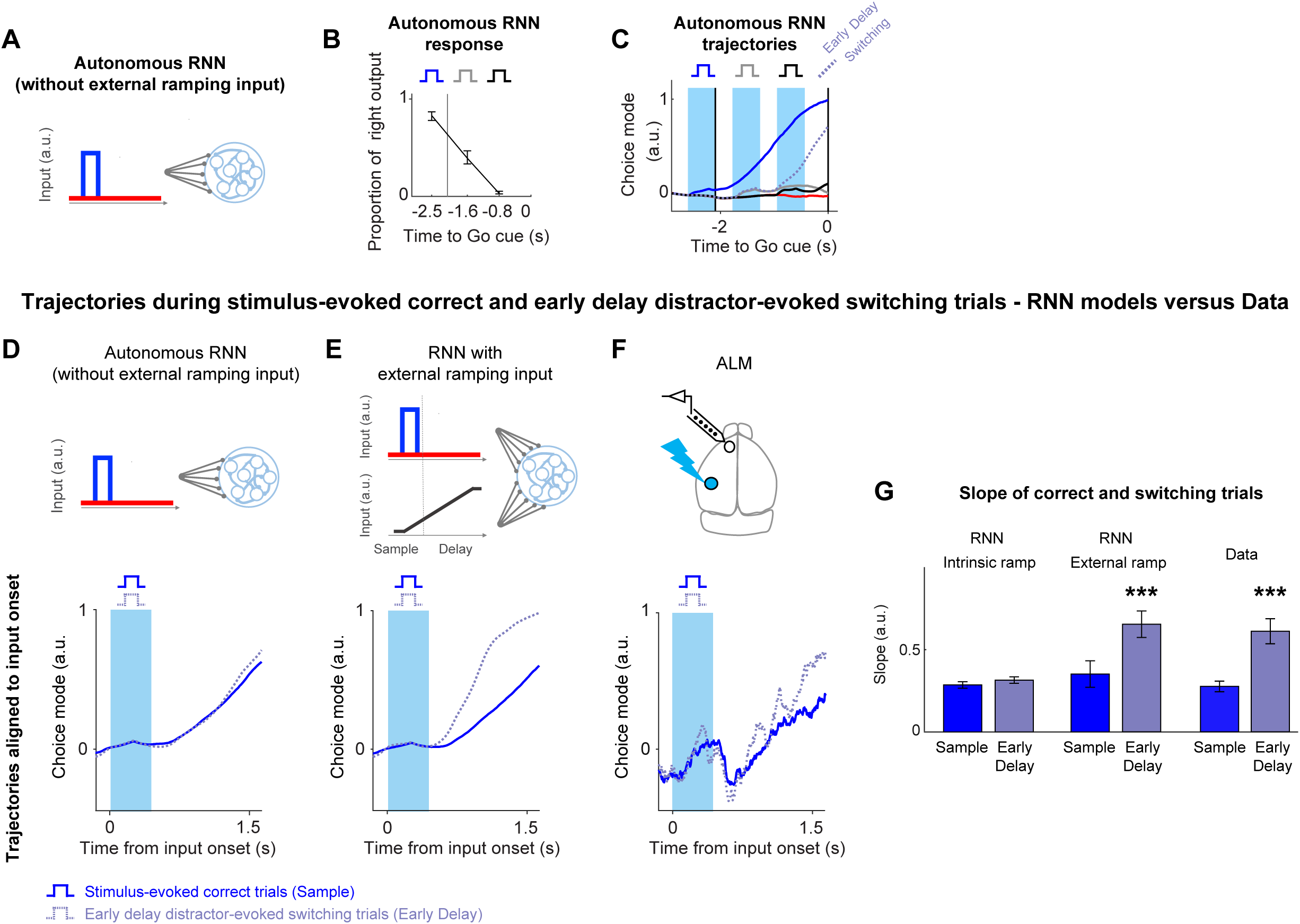
Dynamics of switching trials in RNN models with and without external ramping, comparison with the data. **A-C,** RNN trained without an external ramping input (autonomous RNN). **A,** This model relies on internal recurrent (autonomous) dynamics to reproduce the slow ramping observed in the data. **B-C**, Labels as in Fig. 2C-D. **B,** Proportion of right outputs generated by autonomous RNN. The proportion of lick-right distractor-evoked output trials was similar to those observed in the data and in the RNN with external ramping, implying that the autonomous RNN also displayed a form of temporal gating. **C**, RNN activity projected on Choice mode for correct and switching trials evoked by early-delay distractor (dotted gray). **D-F**, Trial-averaged trajectories of stimulus-evoked correct trials (blue) and early-distractor evoked (gray) switching trials, aligned to stimulus/distractor onset for (**D**) autonomous RNN, (**E**) RNN with external ramping, and (**F**) in experimental data, i.e. left ALM. **G,** Slopes of stimulus-evoked correct trials (blue) and early-distractor-evoked (gray) switching trials trajectories in autonomous RNN, RNN with external ramping, and left ALM. Error bars, mean ± s.e.m. across trials (RNNs) or across sessions (data). Early-delay switching trajectories in the data were similar to trajectories generated by the RNN model with external ramping input, but not to the ones generated by the autonomous RNN.

**Supplementary Figure 9.**
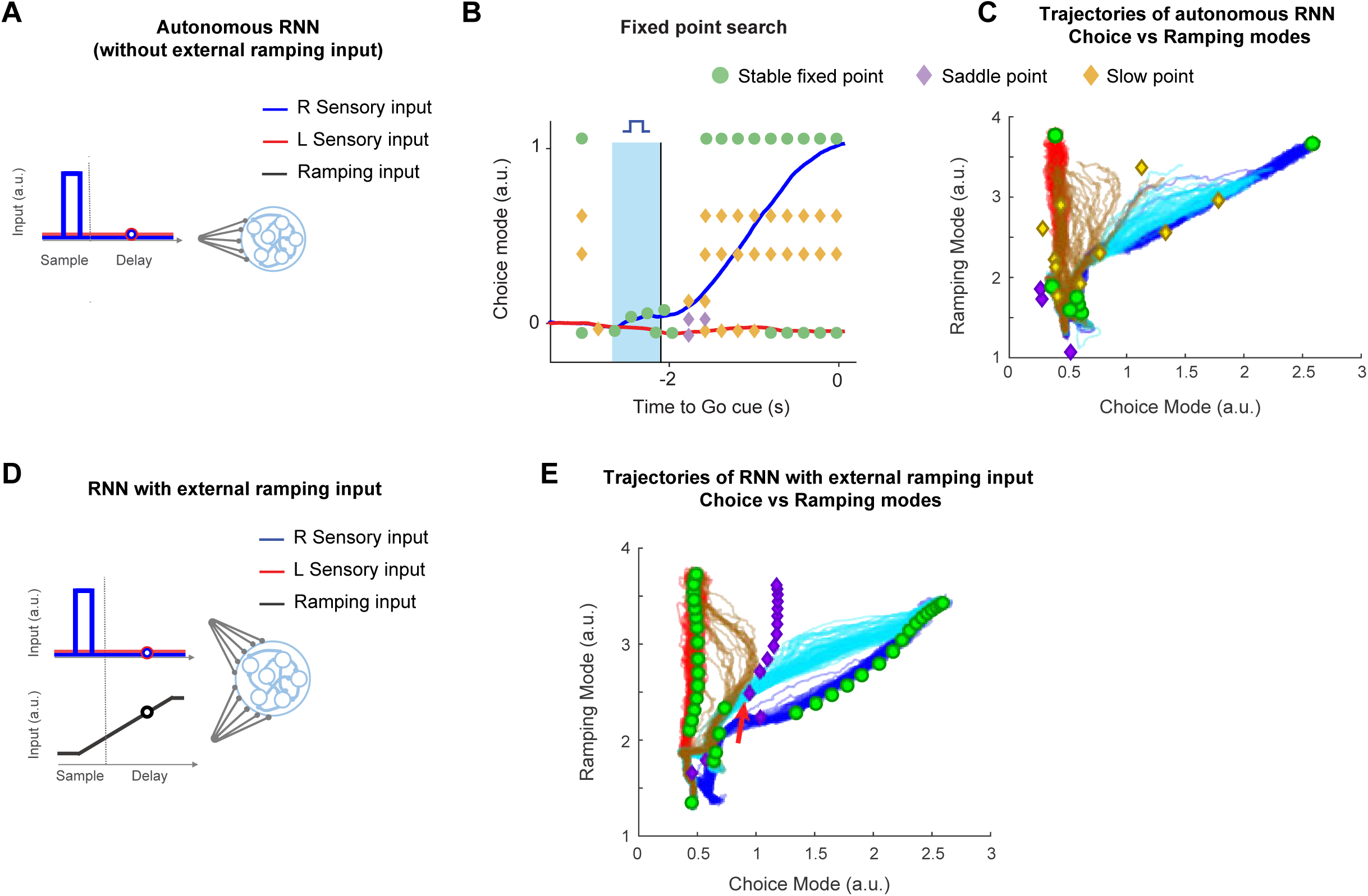
Fixed-point analysis of autonomous RNN and 2-D projections of RNN trajectories. **A**, Autonomous RNN inputs (auditory sample cues are not shown for brevity). Top: transient input during sample epoch. Right trials (blue), left trials (red). Bottom: constant external input (black). For fixed-point search at different time bins the inputs were fixed at values corresponding to each time bin (e.g. t = −0.5 sec, round circles). **B**, Fixed point search for autonomous RNN projected onto Choice mode (same analysis as in Fig. 3E for RNN with external ramping). Slow points of the dynamics (i.e. points were the flow was significantly faster than zero but much slower than single unit dynamics, see Methods) are showed in yellow. Notice that during the delay epoch there are no stable fixed points close to the trajectories (except at the very end), contrary to RNN with external input (**Fig. 3E**). **C**, 2-D projections of neural trajectories on Choice and Ramping modes, and location of fixed point. Shown are projections averaged across correct trials in response to stimulus (blue), no stimulus (red), switching trials evoked by early-delay distractor (cyan) and robust trials evoked by early-delay distractor (brown). Stable fixed points are shown as green circles, saddle points as purple diamonds, and slow points as yellow diamonds. Slow ramping dynamics is achieved by the autonomous RNN by creating slow points (yellow diamonds) along the desired lick-right and lick-left trajectories. Two stable fixed points are located at the end of lick-right and lick-left trajectories. **D**, Inputs of RNN with external ramp (auditory cues are not shown for brevity). Top: transient input during sample epoch. Right trials (blue), left trials (red). Bottom: external input with a linearly ramping component in both right and left trials (black). As for the autonomous RNN, for fixed-point search at different time bins the inputs were fixed at values corresponding to each time bin (e.g. t = −0.5 sec, round circles). **E**, Same as in C, for RNN trained with external ramping input. Correct and switching trajectories separate in the vicinity of the first saddle point (red arrow). We did not find slow points in the RNN trained with external ramping.

**Supplementary Figure 10.**
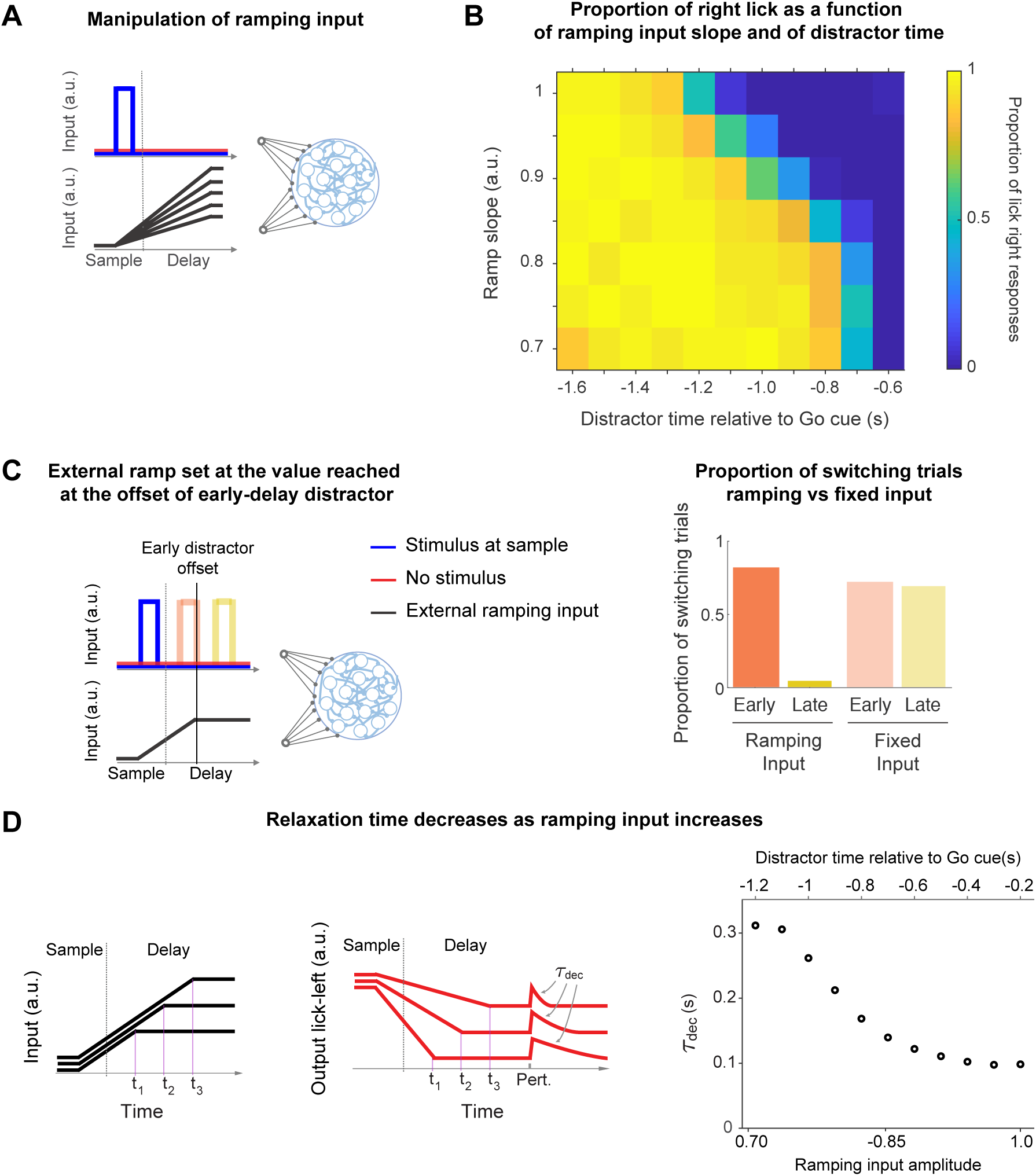
External ramping input level determines robustness to distractors. **A**, RNN model with external ramping input of various strength. **B,** Proportion of distractor-evoked switching trials as a function of the time at which the distractor was delivered and of ramping input strength. Note that i) weaker ramping input induce more errors and ii) distractors delivered late in the delay are more efficiently gated. **C,** Assessing the contribution of external ramping input to temporal gating of distractors, in the RNN trained in presence of ramping input. **C,** left panel: To dissociate the impact on RNN dynamics of the external ramping input, which progressively increased during sample and delay epochs, from the effects of passage of time only, we clamped the external ramping input at the level it reached at the offset of the early-delay distractor (t = −1.2 s, black vertical line). Right panel: Proportion of switching trials for early- and late-delay distractors: unlike the case of Fig. 3A, where the external input kept ramping throughout the delay epoch, the proportion of switching trials evoked by late-delay distractors when the ramping input was clamped was similar to that of early-delay distractors. This suggests that temporal gating is regulated by the external ramp level and not by the passage of time *per se*. **D**, Relaxation time constants of RNN with external input as a function of time at which the perturbation was delivered. In this test, the ramping input was set to the value it had reached at time _*#*_ in the delay (left panel), and, for each ramping input level, a perturbation of amplitude 0.2 a.u. along the Choice mode was delivered to the RNN during lick-left trials (middle panel). The relaxation time constant decreased as ramping slope increased (right panel), suggesting a deepening of the lick-left attraction basin with increased ramping (which corresponds, equivalently, to later times in the delay).

**Supplementary Figure 11.**
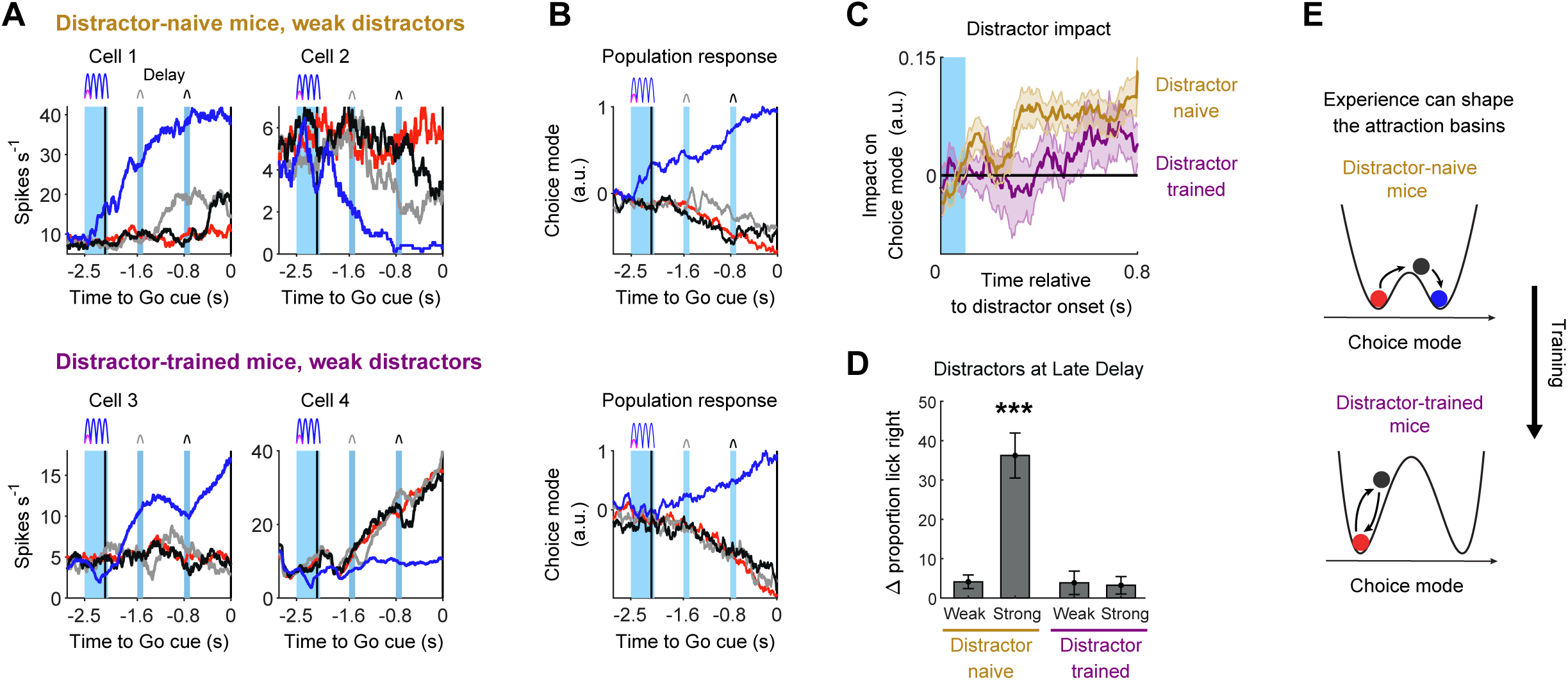
Distractor-impact on persistent activity in ALM is learning-dependent. Spike rate modulations in left ALM in distractor-naive versus distractor-trained mice, in response to weak-distractors computed using correct trials. **A,** Spike rates of example cells and **B,** Activity along the Choice mode in the presence of weak distractors in distractor-naive (top) and distractor trained mice (bottom). Lick-left trajectory without stimulation (red), and with weak distractors during early-delay (gray), or late-delay (black). Lick-right trajectory during sample-epoch stimulation (blue). Note that weak distractors had a persistent effect in ALM of distractor-naive mice. Specifically, distractors shifted the activity from lick-left trajectory towards lick-right trajectory. This was evident from shifts in projections on Choice mode (B) and from activity of individual cells that contributed to this mode (A). In contrast, in distractor-trained mice, the effect of distractors was transient. **C,** Impact of weak distractors on Choice mode in distractor-naive (blue) and distractor-trained mice (green). Trajectories were aligned to the onset of each distractor. Data is shown as average across distractors with different onset times and sessions ± s.e.m. (shaded). In distractor-naive mice the effect of distractor persisted for at least 0.8 s (the time interval from the late-delay distractor onset and the Go cue), whereas in distractor-trained mice the activity recovered to the unperturbed trajectory. **D,** Change in proportion of lick-right responses on late-delay distractor trials (weak and strong distractors) relative to no-stimulus trials, in distractor-naive versus distractor trained mice. Error bars, mean ± s.e.m. across sessions; *** *P* < 0.001 by paired Student t-test (distractor trials versus non-stimulus trials). **E,** Schematics of putative attraction basins in distractor-naive (top) and distractor-trained (bottom) mice. Shallow basin of attraction in distractor-mice mice allowed sufficiently strong stimuli to switch the neural activity from one basin to another.

## References

1. J. M. Fuster, G. E. Alexander, Neuron Activity Related to Short-Term Memory. Science. 173, 652–654 (1971).

2. K. Kubota, H. Niki, Prefrontal cortical unit activity and delayed alternation performance in monkeys. Journal of Neurophysiology. 34, 337–347 (1971).

3. J. Tanji, E. V. Evarts, Anticipatory activity of motor cortex neurons in relation to direction of an intended movement. Journal of Neurophysiology. 39, 1062–1068 (1976).

4. Y. Miyashita, H. S. Chang, Neuronal correlate of pictorial short-term memory in the primate temporal cortexYasushi Miyashita. Nature. 331, 68–70 (1988).

5. I. Fried, R. Mukamel, G. Kreiman, Internally Generated Preactivation of Single Neurons in Human Medial Frontal Cortex Predicts Volition. Neuron. 69, 548–562 (2011).

6. J. C. Erlich, M. Bialek, C. D. Brody, A Cortical Substrate for Memory-Guided Orienting in the Rat. Neuron. 72, 330–343 (2011).

7. A. K. Churchland, R. Kiani, M. N. Shadlen, Decision-making with multiple alternatives. Nature Neuroscience. 11, 693–702 (2008).

8. Z. V. Guo, N. Li, D. Huber, E. Ophir, D. Gutnisky, J. T. Ting, G. Feng, K. Svoboda, Flow of Cortical Activity Underlying a Tactile Decision in Mice. Neuron. 81, 179–194 (2014).

9. M. T. Kaufman, M. M. Churchland, S. I. Ryu, K. V. Shenoy, Cortical activity in the null space: permitting preparation without movement. Nat Neurosci. 17, 440–448 (2014).

10. M. T. Kaufman, M. M. Churchland, S. I. Ryu, K. V. Shenoy, Vacillation, indecision and hesitation in moment-by-moment decoding of monkey motor cortex. eLife. 4, e04677 (2015).

11. N. Li, K. Daie, K. Svoboda, S. Druckmann, Robust neuronal dynamics in premotor cortex during motor planning. Nature. 532, 459–464 (2016).

12. J. A. Michaels, B. Dann, R. W. Intveld, H. Scherberger, Neural Dynamics of Variable Grasp-Movement Preparation in the Macaque Frontoparietal Network. J. Neurosci. 38, 5759–5773 (2018).

13. K. Svoboda, N. Li, Neural mechanisms of movement planning: motor cortex and beyond. Current Opinion in Neurobiology. 49, 33–41 (2018).

14. H. K. Inagaki, L. Fontolan, S. Romani, K. Svoboda, Discrete attractor dynamics underlies persistent activity in the frontal cortex. Nature. 566, 212 (2019).

15. J. Moran, R. Desimone, Selective attention gates visual processing in the extrastriate cortex. Science. 229, 782–784 (1985).

16. E. Seidemann, E. Zohary, W. T. Newsome, Temporal gating of neural signals during performance of a visual discrimination task. Nature. 394, 72–75 (1998).

17. T. Moore, K. M. Armstrong, Selective gating of visual signals by microstimulation of frontal cortex. Nature. 421, 370–373 (2003).

18. V. Mante, D. Sussillo, K. V. Shenoy, W. T. Newsome, Context-dependent computation by recurrent dynamics in prefrontal cortex. Nature. 503, 78–84 (2013).

19. M. G. Stokes, M. Kusunoki, N. Sigala, H. Nili, D. Gaffan, J. Duncan, Dynamic Coding for Cognitive Control in Prefrontal Cortex. Neuron. 78, 364–375 (2013).

20. S. Tremblay, F. Pieper, A. Sachs, J. Martinez-Trujillo, Attentional Filtering of Visual Information by Neuronal Ensembles in the Primate Lateral Prefrontal Cortex. Neuron. 85, 202–215 (2015).

21. R. D. Wimmer, L. I. Schmitt, T. J. Davidson, M. Nakajima, K. Deisseroth, M. M. Halassa, Thalamic control of sensory selection in divided attention. Nature. 526, 705–709 (2015).

22. D. Xu, Y. Chen, A. M. Delgado, N. C. Hughes, L. Zhang, M. Dong, D. H. O’Connor, A functional cortical network for sensorimotor sequence generation. bioRxiv (2019), doi: 10.1101/783050.

23. Z. Wu, A. Litwin-Kumar, P. Shamash, A. Taylor, R. Axel, M. N. Shadlen, Context-dependent decision making in a premotor circuit. bioRxiv, 757104 (2019).

24. D. Huber, L. Petreanu, N. Ghitani, S. Ranade, T. Hromádka, Z. Mainen, K. Svoboda, Sparse optical microstimulation in barrel cortex drives learned behaviour in freely moving mice. Nature. 451, 61–64 (2008).

25. D. H. O’Connor, S. A. Hires, Z. V. Guo, N. Li, J. Yu, Q.-Q. Sun, D. Huber, K. Svoboda, Neural coding during active somatosensation revealed using illusory touch. Nat Neurosci. 16, 958–965 (2013).

26. M. H. Histed, J. H. R. Maunsell, Cortical neural populations can guide behavior by integrating inputs linearly, independent of synchrony. PNAS. 111, E178–E187 (2014).

27. V. de Lafuente, R. Romo, Neural correlate of subjective sensory experience gradually builds up across cortical areas. PNAS. 103, 14266–14271 (2006).

28. A. Hernández, V. Nácher, R. Luna, A. Zainos, L. Lemus, M. Alvarez, Y. Vázquez, L. Camarillo, R. Romo, Decoding a Perceptual Decision Process across Cortex. Neuron. 66, 300–314 (2010).

29. K. Rajan, C. D. Harvey, D. W. Tank, Recurrent Network Models of Sequence Generation and Memory. Neuron. 90, 128–142 (2016).

30. D. Sussillo, O. Barak, Opening the Black Box: Low-Dimensional Dynamics in High-Dimensional Recurrent Neural Networks. Neural Computation. 25, 626–649 (2012).

31. A. Resulaj, R. Kiani, D. M. Wolpert, M. N. Shadlen, Changes of mind in decision-making. Nature. 461, 263–266 (2009).

32. A. Bollimunta, D. Totten, J. Ditterich, Neural Dynamics of Choice: Single-Trial Analysis of Decision-Related Activity in Parietal Cortex. J. Neurosci. 32, 12684–12701 (2012).

33. R. Kiani, C. J. Cueva, J. B. Reppas, W. T. Newsome, Dynamics of Neural Population Responses in Prefrontal Cortex Indicate Changes of Mind on Single Trials. Current Biology. 24, 1542–1547 (2014).

34. D. Thura, P. Cisek, Deliberation and Commitment in the Premotor and Primary Motor Cortex during Dynamic Decision Making. Neuron. 81, 1401–1416 (2014).

35. M. Suzuki, J. Gottlieb, Distinct neural mechanisms of distractor suppression in the frontal and parietal lobe. Nature Neuroscience. 16, 98–104 (2013).

36. G. M. Ghose, J. H. R. Maunsell, Attentional modulation in visual cortex depends on task timing. Nature. 419, 616–620 (2002).

37. A. Nobre, A. Correa, J. Coull, The hazards of time. Current Opinion in Neurobiology. 17, 465–470 (2007).

38. S. Jaramillo, A. M. Zador, The auditory cortex mediates the perceptual effects of acoustic temporal expectation. Nature Neuroscience. 14, 246–251 (2011).

39. F. Carnevale, V. de Lafuente, R. Romo, O. Barak, N. Parga, Dynamic Control of Response Criterion in Premotor Cortex during Perceptual Detection under Temporal Uncertainty. Neuron. 86, 1067–1077 (2015).

40. F. van Ede, S. R. Chekroud, M. G. Stokes, A. C. Nobre, Decoding the influence of anticipatory states on visual perception in the presence of temporal distractors. Nat Commun. 9, 1–12 (2018).

41. D. Sussillo, L. F. Abbott, Generating Coherent Patterns of Activity from Chaotic Neural Networks. Neuron. 63, 544–557 (2009).

42. J. Wang, D. Narain, E. A. Hosseini, M. Jazayeri, Flexible timing by temporal scaling of cortical responses. Nature Neuroscience, 1 (2017).

43. G. R. Yang, M. R. Joglekar, H. F. Song, W. T. Newsome, X.-J. Wang, Task representations in neural networks trained to perform many cognitive tasks. Nature Neuroscience. 22, 297 (2019).

44. H. S. Seung, How the brain keeps the eyes still. Proceedings of the National Academy of Sciences. 93, 13339–13344 (1996).

45. D. J. Amit, N. Brunel, Model of global spontaneous activity and local structured activity during delay periods in the cerebral cortex. Cereb Cortex. 7, 237–252 (1997).

46. X.-J. Wang, Probabilistic Decision Making by Slow Reverberation in Cortical Circuits. Neuron. 36, 955–968 (2002).

47. C. D. Brody, R. Romo, A. Kepecs, Basic mechanisms for graded persistent activity: discrete attractors, continuous attractors, and dynamic representations. Current Opinion in Neurobiology. 13, 204–211 (2003).

48. C. K. Machens, R. Romo, C. D. Brody, Flexible Control of Mutual Inhibition: A Neural Model of Two-Interval Discrimination. Science. 307, 1121–1124 (2005).

49. S. Lim, M. S. Goldman, Balanced cortical microcircuitry for maintaining information in working memory. Nature Neuroscience. 16, 1306–1314 (2013).

50. O. Barak, M. Tsodyks, Working models of working memory. Current Opinion in Neurobiology. 25, 20–24 (2014).

51. K. Wimmer, D. Q. Nykamp, C. Constantinidis, A. Compte, Bump attractor dynamics in prefrontal cortex explains behavioral precision in spatial working memory. Nat Neurosci. 17, 431–439 (2014).

52. C. D. Kopec, J. C. Erlich, B. W. Brunton, K. Deisseroth, C. D. Brody, Cortical and Subcortical Contributions to Short-Term Memory for Orienting Movements. Neuron. 88, 367–377 (2015).

53. R. Chaudhuri, I. Fiete, Computational principles of memory. Nat Neurosci. 19, 394–403 (2016).

54. J. Kaminski, S. Sullivan, J. M. Chung, I. B. Ross, A. N. Mamelak, U. Rutishauser, Persistently active neurons in human medial frontal and medial temporal lobe support working memory. Nature Neuroscience. 20, 590–601 (2017).

55. E. K. Miller, C. A. Erickson, R. Desimone, Neural Mechanisms of Visual Working Memory in Prefrontal Cortex of the Macaque. J. Neurosci. 16, 5154–5167 (1996).

56. K. Sakai, J. B. Rowe, R. E. Passingham, Active maintenance in prefrontal area 46 creates distractor-resistant memory. Nat Neurosci. 5, 479–484 (2002).

57. S. N. Jacob, A. Nieder, Complementary Roles for Primate Frontal and Parietal Cortex in Guarding Working Memory from Distractor Stimuli. Neuron. 83, 226–237 (2014).

58. R. Kiani, T. D. Hanks, M. N. Shadlen, Bounded Integration in Parietal Cortex Underlies Decisions Even When Viewing Duration Is Dictated by the Environment. J. Neurosci. 28, 3017–3029 (2008).

59. Y. Zuo, M. E. Diamond, Rats Generate Vibrissal Sensory Evidence until Boundary Crossing Triggers a Decision. Current Biology. 29, 1415–1424.e5 (2019).

60. R. Romo, C. D. Brody, A. Hernández, L. Lemus, Neuronal correlates of parametric working memory in the prefrontal cortex. Nature. 399, 470–473 (1999).

61. A. C. Huk, M. N. Shadlen, Neural Activity in Macaque Parietal Cortex Reflects Temporal Integration of Visual Motion Signals during Perceptual Decision Making. J. Neurosci. 25, 10420–10436 (2005).

62. F. Kouneiher, S. Charron, E. Koechlin, Motivation and cognitive control in the human prefrontal cortex. Nature Neuroscience. 12, 939–945 (2009).

63. M. M. Churchland, J. P. Cunningham, M. T. Kaufman, J. D. Foster, P. Nuyujukian, S. I. Ryu, K. V. Shenoy, Neural population dynamics during reaching. Nature. 487, 51–56 (2012).

64. T. D. Hanks, C. D. Kopec, B. W. Brunton, C. A. Duan, J. C. Erlich, C. D. Brody, Distinct relationships of parietal and prefrontal cortices to evidence accumulation. Nature. 520, 220–223 (2015).

65. J. Poort, A. G. Khan, M. Pachitariu, A. Nemri, I. Orsolic, J. Krupic, M. Bauza, M. Sahani, G. B. Keller, T. D. Mrsic-Flogel, S. B. Hofer, Learning Enhances Sensory and Multiple Non-sensory Representations in Primary Visual Cortex. Neuron. 86, 1478–1490 (2015).

66. A. Parthasarathy, R. Herikstad, J. H. Bong, F. S. Medina, C. Libedinsky, S.-C. Yen, Mixed selectivity morphs population codes in prefrontal cortex. Nature Neuroscience. 20, 1770 (2017).

67. A. Akrami, C. D. Kopec, M. E. Diamond, C. D. Brody, Posterior parietal cortex represents sensory history and mediates its effects on behaviour. Nature. 554, 368–372 (2018).

68. R. Hattori, B. Danskin, Z. Babic, N. Mlynaryk, T. Komiyama, Area-Specificity and Plasticity of History-Dependent Value Coding During Learning. Cell. 177, 1858–1872.e15 (2019).

69. P. Janssen, M. N. Shadlen, A representation of the hazard rate of elapsed time in macaque area LIP. Nature Neuroscience. 8, 234–241 (2005).

70. G. Maimon, J. A. Assad, A cognitive signal for the proactive timing of action in macaque LIP. Nature Neuroscience. 9, 948–955 (2006).

71. G. T. Finnerty, M. N. Shadlen, M. Jazayeri, A. C. Nobre, D. V. Buonomano, Time in Cortical Circuits. J. Neurosci. 35, 13912–13916 (2015).

72. F. P. Chabrol, A. Blot, T. D. Mrsic-Flogel, Cerebellar Contribution to Preparatory Activity in Motor Neocortex. Neuron. 103, 506–519.e4 (2019).

73. L. Madisen, T. A. Zwingman, S. M. Sunkin, S. W. Oh, H. A. Zariwala, H. Gu, L. L. Ng, R. D. Palmiter, M. J. Hawrylycz, A. R. Jones, E. S. Lein, H. Zeng, A robust and high-throughput Cre reporting and characterization system for the whole mouse brain. Nat Neurosci. 13, 133–140 (2010).

74. L. Madisen, T. Mao, H. Koch, J. Zhuo, A. Berenyi, S. Fujisawa, Y.-W. A. Hsu, A. J. Garcia, X. Gu, S. Zanella, J. Kidney, H. Gu, Y. Mao, B. M. Hooks, E. S. Boyden, G. Buzsáki, J. M. Ramirez, A. R. Jones, K. Svoboda, X. Han, E. E. Turner, H. Zeng, A toolbox of Cre-dependent optogenetic transgenic mice for light-induced activation and silencing. Nat Neurosci. 15, 793–802 (2012).

75. S. Pluta, A. Naka, J. Veit, G. Telian, L. Yao, R. Hakim, D. Taylor, H. Adesnik, A direct translaminar inhibitory circuit tunes cortical output. Nature Neuroscience. 18, 1631–1640 (2015).

76. D. Yatsenko, J. Reimer, A. S. Ecker, E. Y. Walker, F. Sinz, P. Berens, A. Hoenselaar, R. J. Cotton, A. S. Siapas, A. S. Tolias, DataJoint: managing big scientific data using MATLAB or Python. bioRxiv, 031658 (2015).

77. J. J. Jun, C. Mitelut, C. Lai, S. L. Gratiy, C. A. Anastassiou, T. D. Harris, Real-time spike sorting platform for high-density extracellular probes with ground-truth validation and drift correction. bioRxiv, 101030 (2017).

